# Temperature and nutrients drive eco-phenotypic dynamics in a microbial food web

**DOI:** 10.1101/2021.08.12.456102

**Authors:** Ze-Yi Han, Daniel J. Wieczynski, Andrea Yammine, Jean P. Gibert

**Author notes:** To whom correspondence should be addressed:* Ze-Yi Han, Department of Biology, Duke University, Durham, NC, USA. Data accessibility statement:* All annotated code and data are available at our dedicated repository (https://github.com/ZeYiHan/Nut_Temp_Pheno_Eco) and Zenodo (DOI: 10.5281/zenodo.7474517).

## Abstract

Anthropogenic increases in temperature and nutrient loads will likely impact food web structure and stability. Although their independent effects have been reasonably well studied, their joint effects—particularly on coupled ecological and phenotypic dynamics—remain poorly understood. Here we experimentally manipulated temperature and nutrient levels in microbial food webs and used time-series analysis to quantify the strength of reciprocal effects between ecological and phenotypic dynamics across trophic levels. We found that i) joint –often interactive– effects of temperature and nutrients on ecological dynamics are more common at higher trophic levels, ii) temperature and nutrients interact to shift the relative strength of top-down vs. bottom-up control, and iii) rapid phenotypic change mediates observed ecological responses to changes in temperature and nutrients. Our results uncover how feedbacks between ecological and phenotypic dynamics mediate food web responses to environmental change. This suggests important but previously unknown ways that temperature and nutrients might jointly control the rapid eco-phenotypic feedbacks that determine food web dynamics in a changing world.

## INTRODUCTION

Understanding how rapid global climate change (GCC) will affect the structure and dynamics of communities is a pressing goal of ecology (Karl 2003; Barbour & Gibert 2021). Increasing temperatures associated with GCC influence the metabolism of individuals (Gillooly *et al*. 2002; Brown *et al*. 2004; Clarke 2006), which strengthens species interactions (Barton *et al*. 2009; O’Connor 2009; Gounand *et al*. 2016), alters community structure (Bartley *et al*. 2019; Gibert 2019; Gauzens *et al*. 2020), and affects ecosystem function (Dossena *et al*. 2012). Additionally, increasing nutrient loads from agricultural runoffs can result in eutrophication and destabilize natural communities (Rosenzweig 1971; Fussmann 2000; Hautier *et al*. 2014), often leading to species loss (Hautier *et al*. 2009). Warming and eutrophication can independently and jointly impact food web structure and stability (Malcolm *et al*. 2006; Tabi *et al*. 2019). Counterintuitively, simultaneous increases in temperature and nutrient load can produce outcomes that are qualitatively different from the combined negative effects of each variable on its own (Binzer *et al*. 2012, 2016). These non-additive (interactive) outcomes are still poorly understood, but central to honing our understanding of GCC impacts on food web structure and dynamics in a highly anthropogenized world.

The mechanisms through which warming and increasing nutrient loads independently influence food webs are relatively well understood (McClelland & Valiela 1998; Carlier *et al*. 2008; Gibert 2019), specifically regarding their impacts on the relative strength of bottom-up and top-down effects (Binzer *et al*. 2012, 2016; Shurin *et al*. 2012). For example, warming can increase predation pressure (Kratina *et al*. 2012), thus decreasing resource biomass, while increasing the proportion of top predators (Vasseur & McCann 2005; O’Connor 2009; Shurin *et al*. 2012). Alternatively, warming can increase metabolic demands while reducing conversion efficiency (Barneche *et al*. 2021), leading to predator starvation at high temperatures, loss of top predators (Gibert *et al*. 2022a), and reduced food chain length (Petchey *et al*. 1999). Unlike warming, increasing nutrient loads tend to increase bottom-up effects, resulting in unstable dynamics and species loss (i.e., paradox of enrichment, Rip and McCann, 2011; Rosenzweig, 1971), often leading to top-heavy, unstable food webs (Rip & McCann 2011). Eutrophication resulting from increasing nutrient loads can also change consumer trophic position, leading to changes in species interactions and food web structure (Lee *et al*. 2021).

While warming and nutrients can independently influence food webs, they can also have non-additive (interactive) effects when acting unison (Binzer *et al*. 2012, 2016; Tabi *et al*. 2019), but these are much less well understood. Although temperature increases are typically considered to be destabilizing (Barton *et al*. 2009), at low temperatures, small temperature increases can cause consumer starvation, stabilizing nutrient-induced instabilities (Binzer *et al*. 2012, 2016). However, at high temperatures, increasing nutrient loads can counter warming-induced consumer starvation by increasing carrying capacity and predator attack rates (Binzer *et al*. 2012). Warming also weakens nutrient-induced increases in community biomass (negative interactive effects, Tabi et al., 2019), in turn influencing food web structure (Sentis *et al*. 2014), species richness, and community composition (Wang *et al*. 2016a).

In addition to their effects on entire food webs, temperature and nutrients can both determine the physiology and morphology of organisms (Rosenblatt & Schmitz 2016; Tabi *et al*. 2019). For example, higher temperatures often result in smaller sizes (Atkinson 1995; Atkinson *et al*. 2003), while nutrient enrichment leads to larger organisms (Irwin *et al*. 2006; Marañón *et al*. 2013). Additionally, temperature and nutrients can interactively affect body size: size increases with higher nutrient loads at low temperatures, but decreases at high temperature (Tabi *et al*. 2019). Although body size is often considered a response variable, it also has well-known effects on population growth and species interactions (Fenchel 1974; Savage *et al*. 2004; Ferenc & Sheppard 2020; Gibert *et al*. 2022b), so rapid body size responses to temperature, nutrients, or both, may have consequences for food web structure and dynamics in warmer climates (Brose *et al*. 2012; Bernhardt *et al*. 2018; Wieczynski *et al*. 2021). But it is still unclear how rapid, environmentally-induced shifts in body size might influence ecological dynamics as they unfold.

Here, we address how temperature and nutrients influence feedbacks between species’ ecological dynamics and rapid changes in their body sizes (phenotypic dynamics) in a tractable microbial food web. We describe observed food web and body size responses to temperature and nutrients across trophic levels and also study the mechanisms of these responses. Specifically, we ask: 1) do temperature and nutrients independently or interactively influence ecological dynamics in this microbial food web? 2) Do these effects alter the relative importance of top-down vs. bottom-up control? 3) Do these effects vary across trophic levels? And, 4) does body size just passively respond to temperature and nutrients, as suggested elsewhere (e.g., Binzer et al 2016, Tabi et al 2019), or does this body size response also play a role in determining how the food web itself responds to environmental change? To address these questions, we manipulate nutrient levels and temperature in a microbial food web composed of a complex bacterial community, a bacterivorous protist (*Tetrahymena pyriformis*), and an omnivorous top predator (*Euplotes sp*.), then track changes in the population densities of all organisms and the body sizes of the protists over time. We use time-series analysis to evaluate the relative strength of top-down vs. bottom-up processes across trophic levels as well as whether and how the observed ecological and phenotypic dynamics influence one another across temperature and nutrient treatments. Our results reveal complex but quantifiable temperature and nutrient effects on food web dynamics that vary predictably across trophic levels by altering the relative strength of trait-mediated bottom-up and top-down effects.

## METHODS

### Culture care

*Euplotes sp*. and *Tetrahymena pyriformis* stock cultures were acquired from Carolina Biological Supply (Burlington, NC, USA) and grown in laboratory conditions for a year prior to this experimental work. Both species were kept in autoclaved liquid protist medium (Carolina Biological Supply), supplemented with one autoclaved wheat seed as a carbon source (Altermatt et al 2014). Protists were fed a mixture of pond bacteria collected from an ephemeral pond at Duke Forest (Wilbur/Gate 9 pond, Lat 36.013914, Long −78.979720) and composed of thousands of bacterial species, as described elsewhere (Rocca *et al*. 2022). We maintained all cultures on a 16:8 light:dark cycle at 22°C and 65% humidity in AL-22L2 Percival growth chambers (Perry, IA, USA).

### Experimental design

Microcosms were set up in autoclaved 250mL borosilicate jars filled with 200mL of protist media. We manipulated temperature and nutrient loads by imposing two temperature levels (22°C/25°C) and two nutrient levels (normal protist media concentration plus one wheat seed – i.e., high nutrients–, or half concentration plus half a wheat seed, –i.e., low nutrients, as done in (Gibert *et al*. 2017) in a factorial design with 4 treatments and 6 replicates per treatment. Day/night cycle and humidity levels mimicked rearing conditions and were kept constant. We inoculated 2 ml of bacterial communities from the stock culture, the same used to rear the protists. The intermediate consumer species, *T. pyriformis*, only preys on bacteria and was introduced at a starting density of 37 ind/ml. The omnivorous consumer species, *Euplotes sp*., preys on both bacteria and the intermediate consumer *T. pyriformis* and was introduced at a starting density of 0.24 ind/ml in all microcosms. We recorded the density of both protist species once per day through fluid imaging (FlowCam; Yokogawa Fluid Imaging Technologies, Portland, ME, USA), Monday through Friday, for 16 days. Fluid imaging generates individual cell raster images that were used to quantify changes in protist size (measured as cell area, in μm^2^) over time. Bacteria density was quantified as optical density at a wavelength of 600nm (OD600), using a BioTEK Epoch-2 microplate spectrophotometer (Winooski, VT, USA).

### Statistical analysis

To test for possible effects of temperature and nutrients on ecological dynamics, we fitted Generalized Additive Mixed Models (GAMM) to time series of species density (OD600 for bacteria) and protist body size, across all treatments, using the ‘mgcv’ package v1.8-31 <u>(Gibert</u>*<u>e</u>t <u>al</u>*<u>. 2017)</u> in R v4.0.2 (R Core Team 2020). To control for temporal autocorrelation, we used an Autoregressive Moving Average (ARMA) correlation structure in the GAMMs using the ‘nlme’ R package v3.1-148. To account for repeated sampling within each replicate, we included replicates as a random intercept in the model. We compared models with additive and/or interactive temperature and nutrient effects, as well as different ARMA correlation structures, using AICc (Appendix I Table S1-2). We discarded *T. pyriformis* phenotypic data from days in which fewer than 10 individuals were measured (i.e., after populations collapsed).

### Characterizing ecological dynamics

To better understand which aspects of the food web dynamics were most influenced by temperature and nutrients, we characterized multiple aspects of the observed ecological dynamics of the bacterial community and the protists across treatments. Specifically, we quantified: 1) initial growth rate (day^-1^) as [ln(Nf) − ln(Ni)]/time for early-dynamics data, (up to day 1 for *T. pyriformis &* bacteria and day 8 for *Euplotes sp*., due to much slower growth), 2) maximum density (individual/ml for protists and OD_600_ for bacteria), measured as average density across replicates on the day with the highest average density, 3) the coefficient of variation (CV = standard deviation/mean) of the temporal population dynamics within treatments (typically used as a measure of stability (McCann 2012)), 4) the time to population collapse in days (only *T. pyriformis*), and 5) the time to population peak in days (only *Euplotes sp*.). Last, we calculated the effect sizes of the significant effects of temperature, nutrients, and their interaction using the function eta_squared() in package ‘effectsize’ v0.7 (Ben-Shachar *et al*. 2020).

### Quantifying top-down/bottom-up effects and eco-phenotypic feedbacks

To understand the mechanisms through which temperature and nutrients affected the observed food web dynamics, we quantified the reciprocal effects of ecological dynamics and body size on each other using Convergent Cross Mapping (CCM) (Sugihara *et al*. 2012). The CCM algorithm has now been used multiple times across ecological systems and taxa reliably to estimate the strength of causal effects between variables for which time-series are available (Matsuzaki *et al*. 2018; Wang *et al*. 2018; Rogers *et al*. 2020; Barraquand *et al*. 2021; Doi *et al*. 2021; Abidha *et al*. 2022; Gibert *et al*. 2022b) and we followed this specialized literature to infer causation in our data (see Appendix II Fig. S1-15). In a nutshell, CCM quantifies the strength of causation of one dynamical variable onto another by measuring the extent to which the time series of one variable can reliably estimate the state of another variable (Sugihara *et al*. 2012). The larger the causal effect of X on Y, the better the ability of Y to predict X, as Y contains more information about X (by virtue of being ‘forced’ by X). Meanwhile, a variable X that does not influence a variable Y cannot be predicted from the dynamics of Y, as no information regarding X is contained in Y (Sugihara *et al*. 2012). The CCM algorithm yields a ‘cross-mapping estimation skill’ in the form of a correlation coefficient (ρ) between observed and predicted points in the time-series (Sugihara *et al*. 2012). The larger this number, the larger the dependence of one variable on the other. Whether there is a cause-effect relationship between two dynamical variables, as opposed to simple correlation, further depends on whether the cross-mapping estimation skill increases with the length of the time series used for this estimation (called the ‘library size’). Whenever such an increase is observed, a causal effect of a variable on another one is likely (i.e. convergence; Sugihara et al., 2012).

We used a modified CCM algorithm that allows for replication in the time series through the R package ‘multispatialCCM’ (Version 1.0, Clark *et al*. 2015). The package can be used to detect causality between shorter but highly replicated time series, like ours. Our time series lacked data on days 5, 6 and 12, 13 across all replicates. However, the CCM algorithm does not allow for missing time points. To resolve this issue we interpolated the time series data for each replicate using three methods: linear interpolation, spline interpolation, and smooth spline interpolation using the ‘approx’, ‘spline’, and ‘smooth.spline’ functions in base R (see Appendix 2). To increase the robustness of our inference, we then performed CCM in each of these time series and averaged the estimation skill from our CCM analysis across these three separate interpolated time-series. That said, CCM results based on each independent smoothing technique held qualitatively (Appendix II Fig. 1-12), which corroborates the robustness of our inference. Additionally, we only used CCM results that showed convergence in cross mapping skill (ρ) with increasing library size (indicating causality) to focus only on likely causal effects between species ecological and phenotypic dynamics (Appendix II Fig. 13-15).

**Figure 1.**
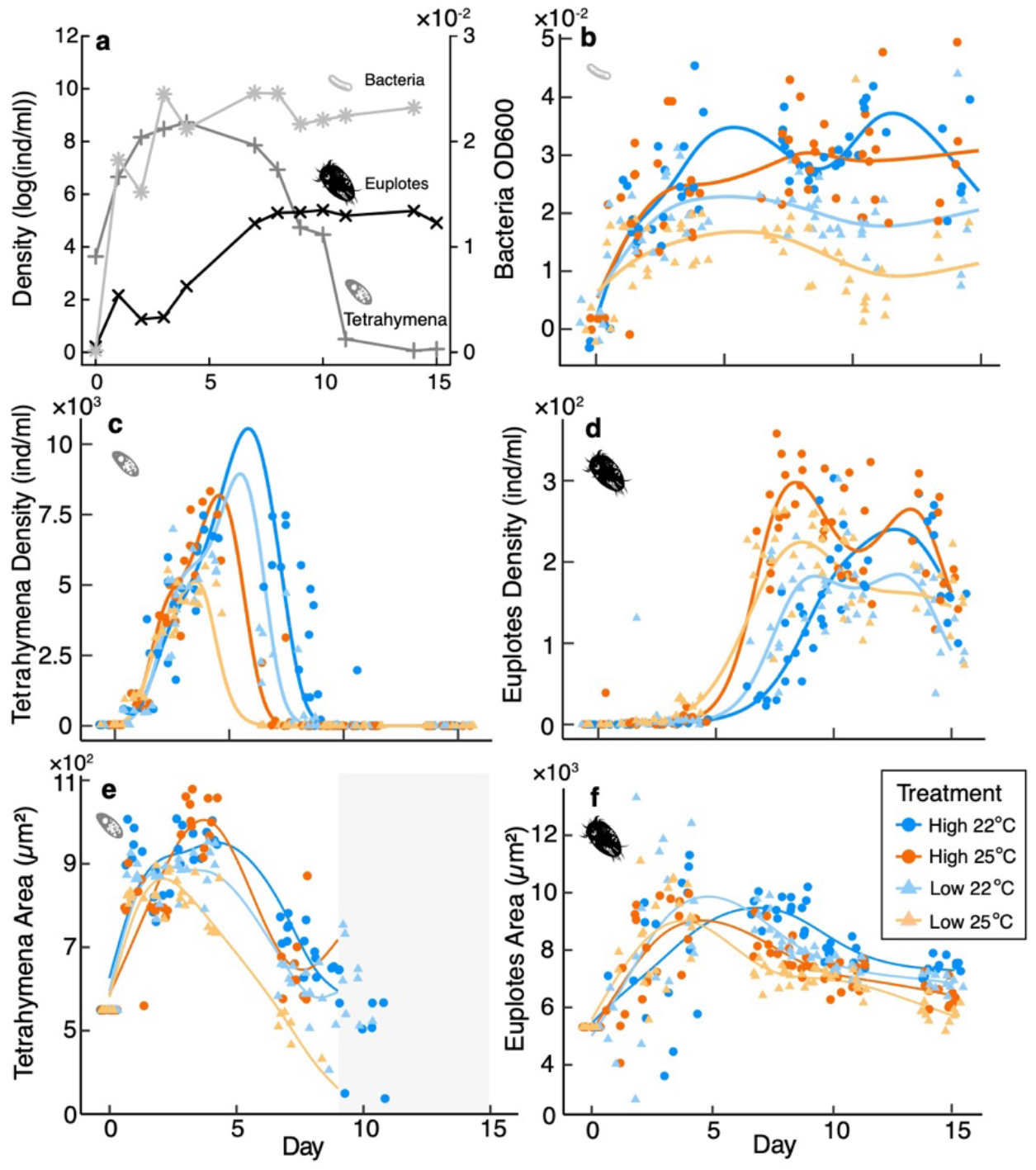
Ecological and phenotypic dynamics within experimental microcosms. (a) Average population densities across treatments. (b-d) Density dynamics of bacteria, *T. pyriformis*, and *Euplotes sp*. (e-f) Changes in the body size (measured as the area of the cell in microm^2) for both protists over time. *T. pyriformis* was effectively extinct by day 9. In (b–f), dots are the empirical measurements and lines are GAMM fits. The greyed area in (e) indicates days for which data were insufficient to estimate trait distributions.

Based on previous literature (Sugihara *et al*. 2012; Matsuzaki *et al*. 2018; Barraquand *et al*. 2021), we interpreted the cross-mapping skill (ρ) as the magnitude of the effect of one variable on another, whenever convergence was present. We calculated this cross-mapping skill between all predator and prey densities, between densities and trait dynamics (except the effect of protist densities on bacterial traits as we lack phenotypic data for the bacteria community), and between protist trait dynamics. The effect of prey density on predator density was thus considered as representing bottom-up control, and the effect of predator density on prey density as top-down control (dubbed “eco-eco” effects, for simplicity). The effects of change in body size on ecological dynamics (density) where dubbed ‘pheno-eco’ effects, and the effects of density on body size dynamics as ‘eco-pheno’ effects. Reciprocal effects of changes in predator and prey protist body sizes were dubbed ‘pheno-pheno’ effects.

## RESULTS

### General Ecological and Phenotypic dynamics

Overall, bacterial density rapidly increased to carrying capacity (Fig. 1a, light grey), the intermediate predator *T. pyriformis* increased rapidly, then decreased (Fig. 1a, dark grey), while the omnivorous predator *Euplotes sp*. increased almost monotonically to carrying capacity (Fig. 1a, black), ultimately resulting in a one protist + bacterial community state. Temperature and nutrients affected all three species and led to significantly different dynamics across treatments (Fig. 1b-d, Appendix I Table S2). The body sizes of both protists changed rapidly over time (Fig. 1e, f) and responded to both temperature and nutrients (Fig. 1e, f, Appendix I Table S2).

### Temperature and nutrient effects changed across trophic levels in systematic ways

We found that both temperature and nutrients significantly and independently affected bacterial dynamics: higher temperature decreased bacterial maximum density, (T_B_ = −0.007, p = 0.004, Fig. 2d), nutrients increased bacterial maximum density (N_B_= 0.014, p = 5.09×10^−6^, Fig. 2d), and nutrients also increased bacterial population density CV (N_B_ = 0.2, p = 8.6×10^−5^, Fig. 2g).

**Figure 2.**
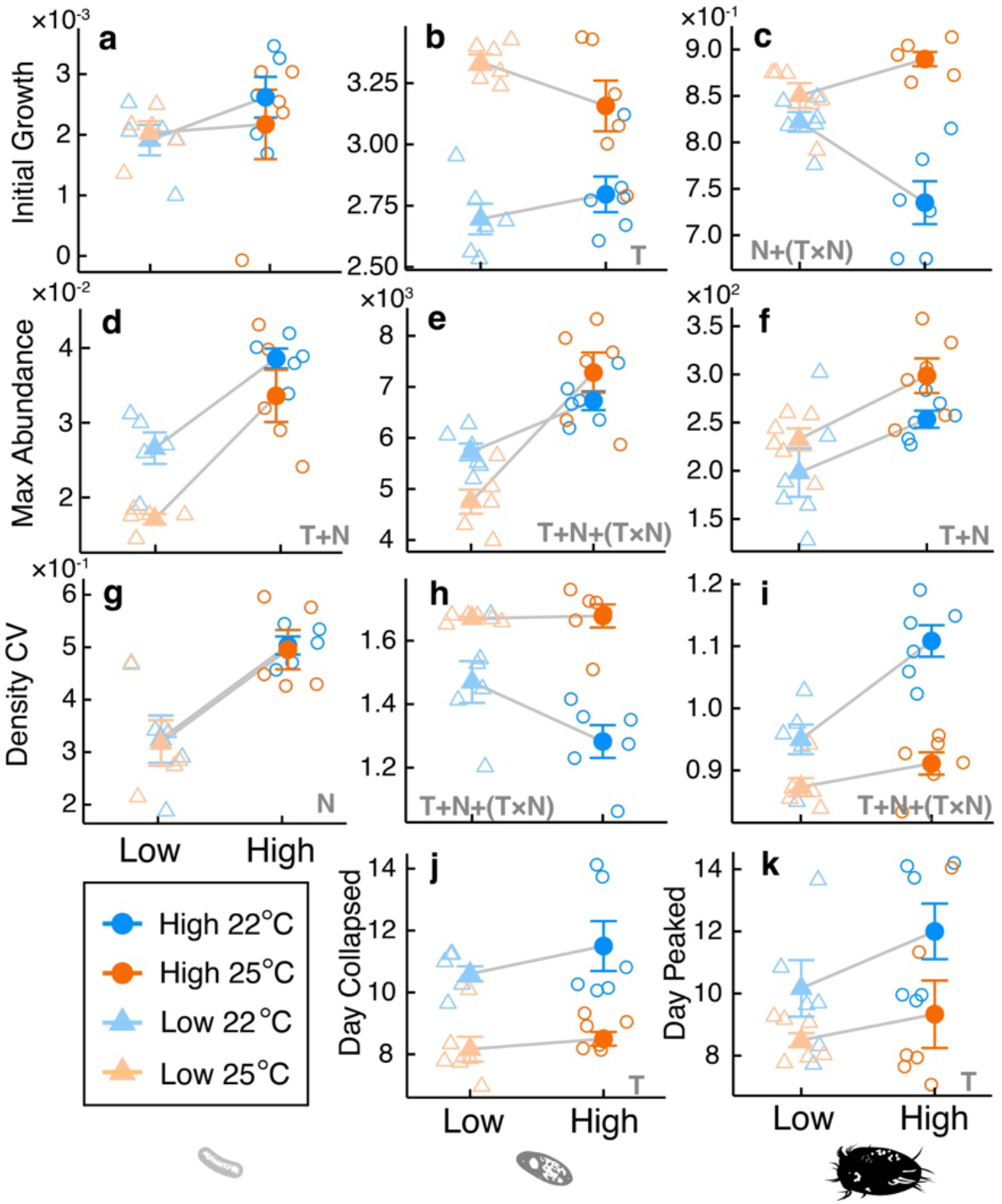
Additive and interactive effects of temperature and nutrients on descriptors of population dynamics. Open circles and triangles represent data points from each replicate. Solid shapes represent means across 6 replicates. Grey letters T, N, and (T×N) represent statistically significant effects from temperature, nutrients, and interactive effects between temperature and nutrients, respectively.

Temperature independently and solely affected *T. pyriformis* initial growth rate and time to population collapse, while temperature and nutrients independently and interactively affected its maximum density and density CV. Temperature strongly increased initial growth rate in *T. pyriformis* (T = 0.50 p = 1.16×10^−6^, Fig. 2b) and accelerated the time to collapse (T = −2.8, p = 1.6×10^−5^; Fig. 2j). The maximum density of *T. pyriformis* increased with nutrients across temperature but only at high nutrient levels, while at low nutrient levels, higher temperature decreased *T. pyriformis* maximum density (T = −975.3, p < 0.02; N = 1002.2, p = 0.01; N×T = 1526.7, p < 0.01, Fig. 2e). The density CV of *T. pyriformis* was also interactively influenced by nutrients and temperature such that at high temperature, *T. pyriformis* CV was high across nutrient levels, but at low temperatures, *T. pyriformis* CV decreased with increasing nutrients (T = 0.2, p < 0.006; N = −0.19, p < 0.01; N×T = 0.2, p = 0.045, Fig. 2h).

Temperature and nutrient levels independently influenced the maximum density and time to the population peak of *Euplotes sp*. and interactively affected its initial growth rate and density CV. Both temperature and nutrients increased the maximum density of *Euplotes sp*. (T = 39.83, p < 0.03; N = 60.67, p < 0.002, Fig. 2f). Temperature alone accelerated *Euplotes* time to peak density (T = −2.2, p < 0.02; Fig. 2k). Higher initial growth rates were observed at higher temperatures, while higher nutrient levels led to large decreases in initial growth rates at low temperatures but a small increase in initial growth rates at high temperatures (N = −0.09, p = 5×10^−4^, N×T = 0.13, p < 10^−3^, Fig. 2c). In general, temperature decreased the density CV of *Euplotes sp*. while higher nutrients increased its CV, more so in lower temperature than in higher temperature (T = −0.08, p < 0.02, N_E_ = 0.16, p = 2.9×10^−5^, N×T = −0.12, p < 0.01, Fig. 2i). Additional model stats can be found in Appendix I Table S3.

We found significant temperature and nutrient interactions only among the predators, with the total effect size of the interaction terms increasing with species trophic level (Fig. 3a). Bacterial population dynamics were mainly influenced by nutrients and the intermediate predator received the most temperature effects (Fig. 3a). These results can be better understood by decomposed total effect sizes of nutrients and temperatures on initial growth rates, maximum density, and density CV. We found no interactive effects of nutrients and temperature on bacteria initial growth rates, but increasingly complex temperature and nutrient effects on species initial growth at higher trophic levels (Fig. 3b). Interestingly, we also found decreasing effect sizes of temperature, nutrients, and their interaction on species maximum density at higher trophic levels (Fig. 3b). However, increasing effect sizes of these effects with increasing trophic level were observed for species density CV (Fig. 3d). These results imply that the maximum density of species at higher trophic levels was less affected by the changes in temperature and nutrients, while the stability of those populations were more sensitive to such changes (Fig. 3).

**Fig. 3.**
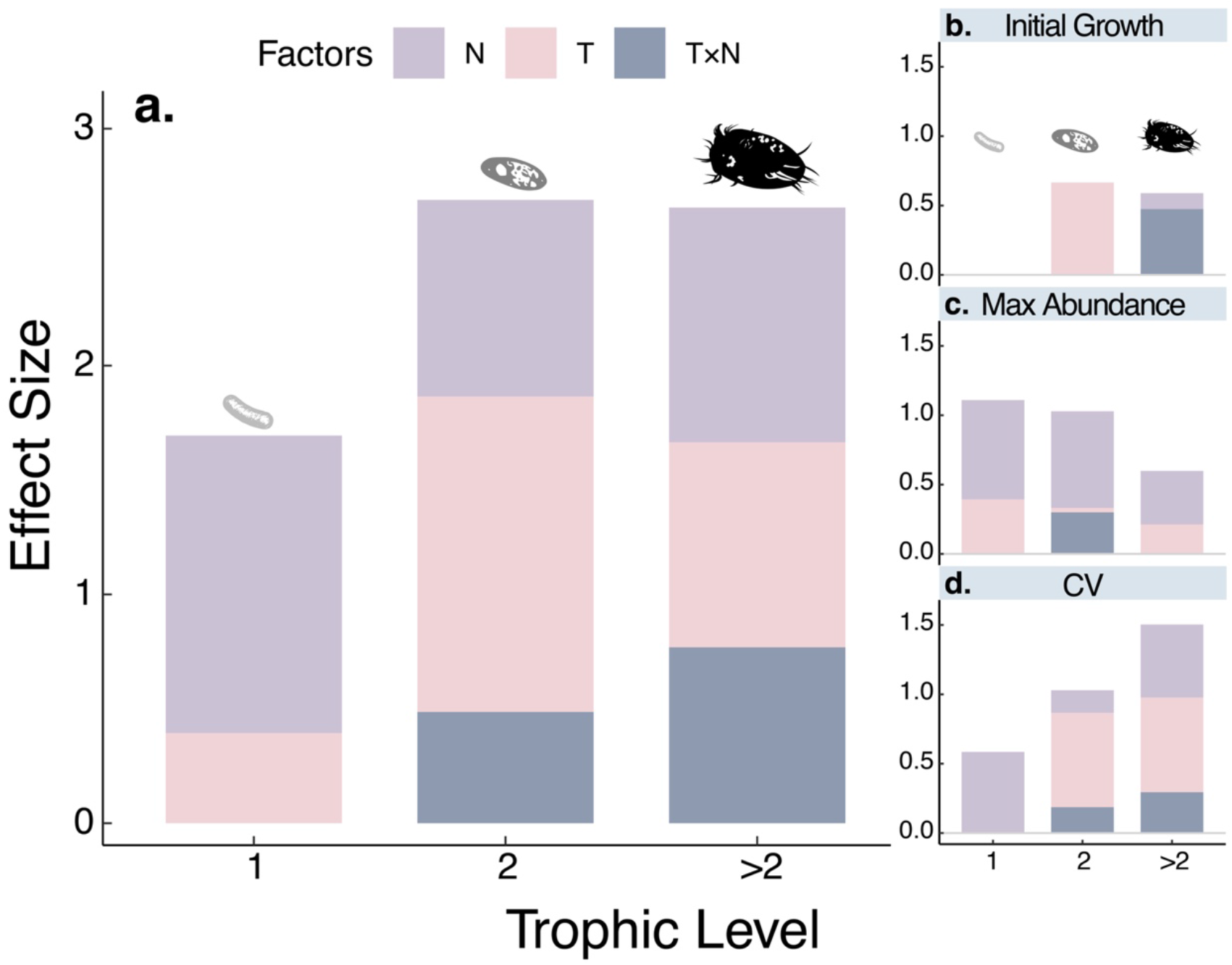
The effect sizes of significant temperature and nutrient effects on ecological dynamics (i.e., initial growth, maximum density, and density CV). T, N, and T×N denote the effects of temperature, nutrients, and interactive effects of temperature and nutrients, respectively.

### Temperature and nutrients influence top-down and bottom-up effects

Results from CCM analyses indicated that the relative strength of bottom-up and top-down effects changed across treatments (Fig. 4a). Top-down and bottom-up effects between the two protists were slightly stronger in the warmer temperature (Fig. 4a, yellow lines). Importantly, temperature and nutrients showed interactive effects on both top-down and bottom-up effects (Fig. 4a). For example, while the top-down effects of protists on bacteria (Fig.4a, green and purple dashed lines) were generally stronger than the bottom-up effects of the bacteria on the protists (Fig. 4a green and purple solid lines), temperature and nutrients interactively –but differentially– shifted the magnitude of the top-down effects of the two protist predators on bacteria (Fig.4a, open dots and dashed lines). Indeed, the top-down effect of *T. pyriformis* on bacteria decreased in the high-nutrient treatment, but higher temperature strengthened this effect (Fig. 4a purple dashed line). Meanwhile, high nutrients increased the top-down effect of *Euplotes sp*. on bacteria in the lower temperature but decreased it at higher temperature (Fig. 4a green dashed line).

**Figure 4.**
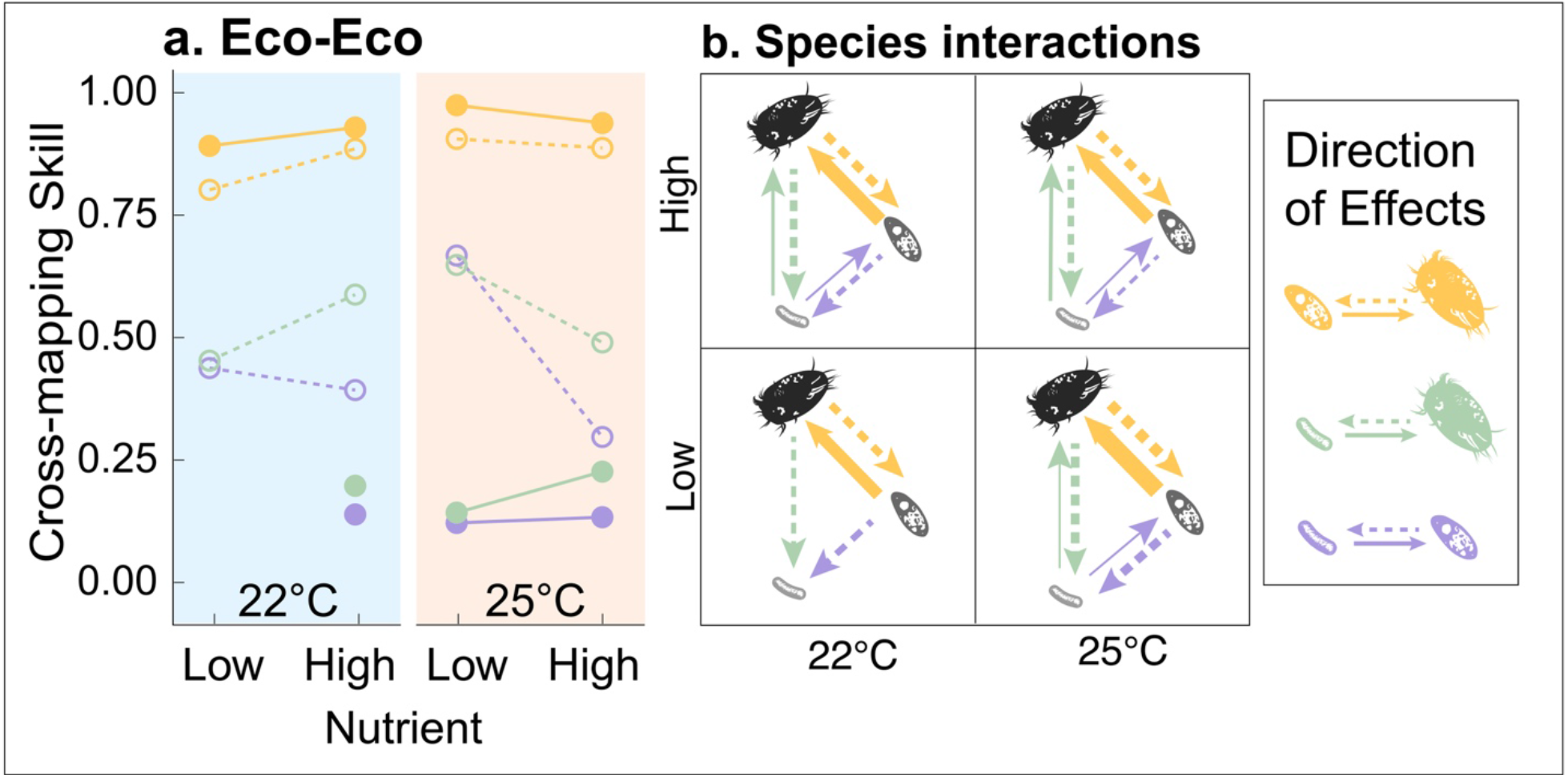
Cross-mapping skill (ρ) of species ecological dynamics across treatments for each species pair. (a) Shows CCM estimation skill of one species’ effect on another. Solid circles and solid lines represent bottom-up effects and open circles and dashed lines represent top-down effects. (b) Shows all causal species interactions for all species considered. Line widths are proportional to the magnitude of the CCM skill while the size of the arrowheads is fixed.

The strength of species interactions in the food web, overall, also changed according to shifts in the strength of top-down and bottom-up control between species pairs (Figure 4b). To visualize these changes, we present CCM cross-mapping skill value at the largest library size as links between species pairs (Fig. 4b). At low nutrient levels, higher temperature led to a larger number of stronger species interactions (Fig. 4b, bottom left and right) while at high nutrient levels, interaction strengths were generally stronger and temperature only had a small effect (Fig. 4b, top left and right). Low nutrients and high temperatures led to the strongest interactions among all three species (Fig. 4b, bottom right).

### Temperature and nutrients altered the reciprocal effects of ecological and phenotypic dynamics

Protist size not only responded to temperature and nutrients, but also played an important role in determining the effects of temperature and nutrients on ecological dynamics (Fig. 5a-c). The bidirectional effects between the body-size dynamics of *Euplotes sp*. and the ecological dynamics of *T. pyriformis* were the strongest overall across treatments (Fig. 5a-c). The body size dynamics of the omnivorous predator, *Euplotes sp*., but not those of *T. pyriformis*, had relatively strong causal effects on the ecological dynamics of all species, including its own ecological dynamics (Fig. 5b). Specifically, at low temperature, increasing nutrient level strengthened the effect of *Euplotes sp*. phenotypic dynamics on its own ecological dynamics and those of *T. pyriformis*, but weakened these effects on the ecological dynamics of bacteria (Fig. 5b). Yet, at the warmer temperature, increasing nutrient levels weakened the effects of *Euplotes sp*. phenotypic dynamics on the ecological dynamics of all species, including itself (Fig. 5b), indicating that plastic changes in top predator body size can mediate how food web dynamics respond to temperature and nutrients.

**Figure 5.**
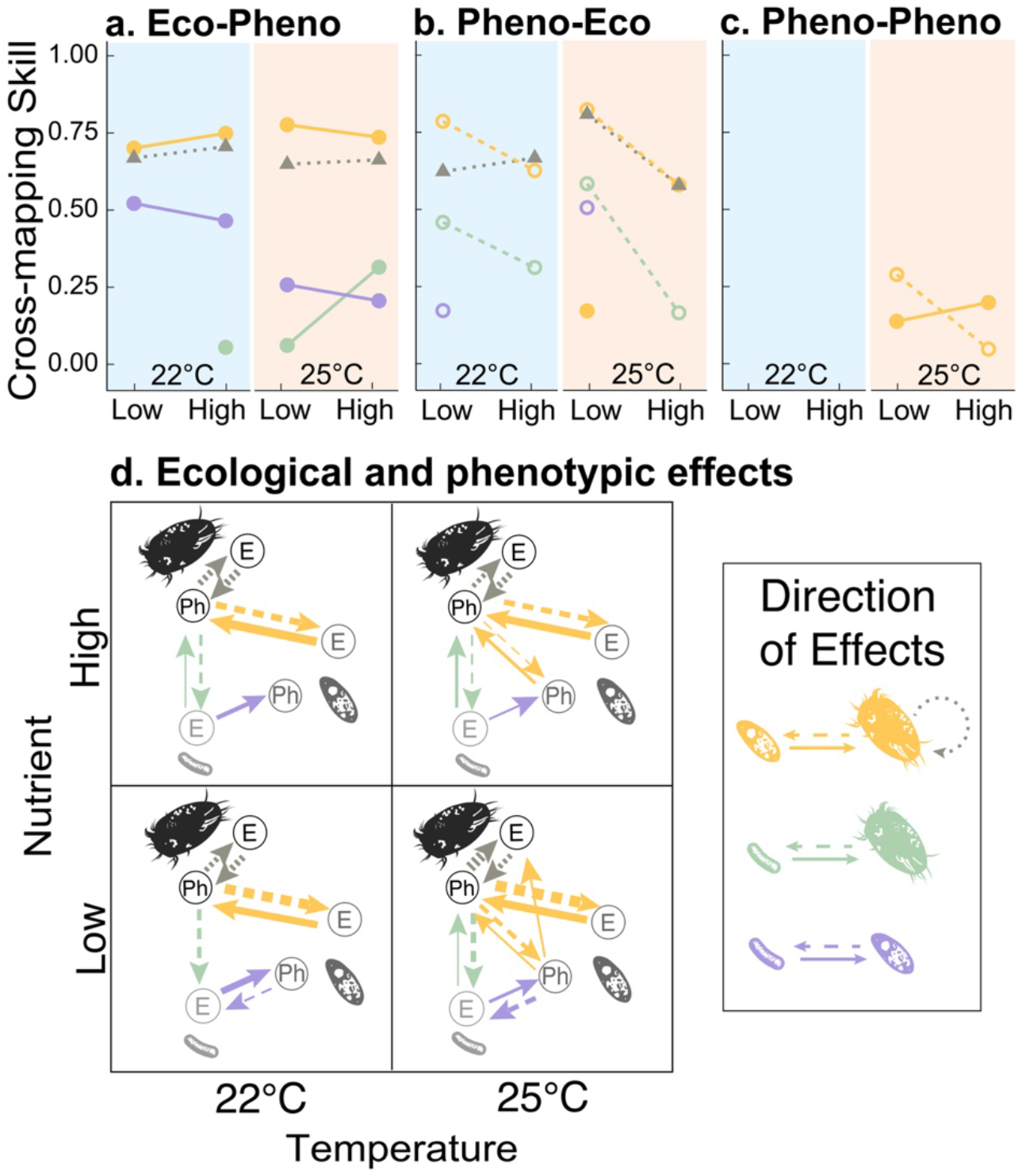
Interactions between phenotypic (Ph) and ecological (E) dynamics. (a-c) Show CCM estimation skill of density on traits (eco-pheno), traits on density (pheno-eco), and trait on trait (pheno-pheno). Circles represent CCM estimation skill of one species’ effect on another species, while triangles represent intraspecific effects. Dashed and solid lines represent top-down and bottom-up effects respectively. Grey dotted lines represent intraspecific effects of *Euplotes sp*. phenotypic dynamics on its own ecological dynamics, and vice versa. In (d), line widths are proportional to the CCM skill while the size of the arrowheads is fixed. Color code as in Fig. 4a.

Trait on trait effects among the two predators (Fig. 5c, pheno-pheno) were weaker than their pheno-eco or eco-pheno counterparts. Interestingly, pheno-pheno causal effects were only observed at the higher temperature (Fig. 5c). Increasing nutrient levels increased the pheno-pheno effects of *T. pyriformis* on *Euplotes sp*. but decreased those of *Euplotes sp*. on *T. pyriformis* (Fig.5c).

Overall, we also found a larger number of stronger effects between ecological and phenotypic dynamics at higher temperatures, especially in the low nutrient treatment (Fig. 5d, bottom right), consistent with results from the top-down vs bottom-up effects (Fig 4b). Additionally, changes in body size, especially those of the omnivorous predator, seem to mediate the effect of temperature and nutrients on ecological dynamics (Fig. 5d).

## DISCUSSION

Our results reveal complex but quantifiable and systematic effects of temperature and nutrients on the ecological and phenotypic dynamics of a microbial food web. We show that temperature and nutrients can each independently influence different aspects of food web dynamics (Fig. 1, 2), while their joint effects get increasingly complex at higher trophic levels (Fig. 2, 3). We also found that changes in the relative strength of top-down and bottom-up effects likely drive observed responses to temperature and nutrients (Fig. 4, 5) and that rapid changes in body size mediate these effects (Fig. 4, 5). Stronger species interactions at low nutrient levels and high temperature coupled with phenotypic dynamics having more and stronger effects in both ecological and phenotypic change at low nutrients and high temperature, suggest that phenotypic change may mediate the temperature response of ecological dynamics, perhaps strengthening species interactions (Fig 4, 5). These results therefore suggest that body size not only responds to shifts in environmental conditions, but also plays a role in determining ecological responses to such shifts (Fig. 5). Evaluating feedbacks between ecological and phenotypic dynamics may therefore be integral to understanding food web responses to environmental change.

### Increasingly complex effects of temperature and nutrients at higher trophic levels

Our results reveal the pervasive effects of temperature and nutrients within a microbial food web, but also show how these effects are more numerous and increasingly complex (i.e., larger effect sizes of temperature-nutrient interactions) at higher trophic levels (Fig. 2, 3). Because energy enters at the bottom of a food web, basal species may be more strongly influenced by direct effects of nutrients, while species at higher trophic levels may be more strongly affected by a combination of temperature and nutrient treatments (Petchey *et al*. 1999; Voigt *et al*. 2003). Additionally, <u>1/5/23 5:11:00 PM</u>the effects of temperature and nutrients on dietary preferences, species interactions, and foraging behavior, may further explain why these interactive effects are stronger and more numerous among consumer species (Fig. 1d; Fig. 2c). Indeed, as temperature and nutrients change species interactions and foraging behaviors (Sentis *et al*. 2014), omnivorous consumers may shift diets between basal and intermediate resources (Sentis *et al*. 2014), leading to climate-driven food web rewiring (Bartley *et al*. 2019; Barbour & Gibert 2021). This dietary shift might also affect how much energy top consumers receive from basal species vs intermediate consumers, and therefore, how temperature and nutrients indirectly influence top predator dynamics. In such case, causal effects of both environmental stressors are likely transitive (e.g. as temperature affects bacteria, and because *Euplotes sp*. preys on bacteria, the temperature effects on bacteria and *Euplotes sp*. are indirectly but casually linked, Sugihara *et al*. 2012).

This transitivity of causality (where omnivorous consumers are indirectly affected by temperature and nutrients that first acted on the basal and intermediate species) could also explain why the population variability (CV) of the top (omnivorous) predator is more often jointly influenced by temperature and nutrients than at lower trophic levels. Moreover, maximum density is more likely to reflect changes in nutrient availability, and these effects should wane across trophic levels, as energy is lost to energy conversion between consumers and prey across trophic levels. And this could explain the decrease of temperature and nutrient effects on species maximum density at higher trophic levels. Whether the joint effects of temperature and nutrients change in more natural food webs as reported here, however, is not known, but is a promising avenue for future research.

### Top-down control, bottom-up control, and food web stability

Consistent with our results (Fig. 2d), theory predicts that higher nutrient loads should increase energy flux in food webs, through increases in basal species density (Rip & McCann 2011; Shurin *et al*. 2012). Theory also predicts that increasing energy flux can be destabilizing, leading to increasing oscillations in density (Rosenzweig 1971; Rip & McCann 2011), but increasing temperatures should stabilize oscillations by weakening top-down effects (Vasseur & McCann 2005; Binzer *et al*. 2016; Tabi *et al*. 2019). We found that the interactive effects of temperature and nutrients have divergent impacts on stability—with temperature stabilizing, and nutrients destabilizing *Euplotes sp*. densities, but the opposite being true for *T. pyriformis* (Fig. 2h-i). Temperature and nutrients had different but interactive effects on top-down controls by the two predators even though bottom-up effects remained unaffected by the treatments (Fig. 4a). These results thus indicate that changes in the strength of top-down effects between basal resources and consumers—instead of bottom-up effects—could be the dominant mechanism through which higher temperatures may stabilize instabilities caused by nutrients (Binzer *et al*. 2012, 2016).

We also observed that top-down control on bacteria by both consumers was much stronger than bottom-up effects across treatments (Fig. 4a) while the top-down and bottom-up controls are both strong and tightly coupled between the two protists predators. The dominant top-down effects on the bacterial community could be explained by the short generation time of all species in the system and the fast turnover rate of the bacteria relative to the protists. In addition to the potential stabilizing effects of top-down control, these results also indicate that top-down control on basal resources might have stronger effects in systems with high turnover rates, such as aquatic systems.

### Phenotypes mediate temperature and nutrient effects on food web dynamics

Our results indicate that phenotypic dynamics play a larger role in mediating environmental impacts on food web dynamics than previously thought (Fig. 4, 5). Previous studies showed that predator-prey body size ratios significantly influence temperature and nutrients effects (Binzer *et al*. 2012; Gibert & DeLong 2014) and that body size responds to changes in nutrients, temperature, and ecological dynamics in specific ways (Tabi *et al*. 2019). A recent study showed that the body size of a species can affect its own ecological dynamics (Gibert *et al*. 2022b), which we have also shown here as tight coupling between the phenotypic and ecological dynamics of *Euplotes sp*. (Fig. 5). Our results support—but also extend—previous findings by showing that 1) phenotypic effects on ecological dynamics can be strong, especially among top predators, 2) changes in the prey ecological dynamics may strongly drive changes in predator size, and 3) interactions involving phenotypic dynamics (eco-pheno, pheno-eco, or pheno-pheno) vary across environmental conditions (Fig. 4b-d, Fig. 5).

### Caveats

One caveat of this study is that we only tracked the dynamics of the bacterial community as a whole, as we lack information on how each individual species responded to temperature or nutrients. Recent work has shown that the composition of bacterial communities changes under joint temperature and nutrient loads, and that predation by protists mediates these responses (Thurman *et al*. 2010; Rocca *et al*. 2022). Coupling an experiment like ours with 16S amplicon sequencing to keep track of bacterial dynamics would thus be an exciting avenue for future research that should deepen the understanding generated by our current study as to how microbial food webs will respond to rapid environmental change.

Another caveat is that, despite uniquely detailed, long, and well-replicated time-series, our time-series have a few small gaps due to lack of sampling on weekends. To address this issue, we interpolated these missing datapoints using three different methods, but, despite being generally robust, some variation remained between these results and the CCM inference of species interactions (Appendix II Figure 1-12). All three methods showed that bottom-up and top-down effects between the protist predators were the strongest and remained strong across all treatments. Moreover, top-down controls from both protists predators to bacteria were always stronger than bottom-up effects (Appendix II Figure 13-15 Eco-Eco panels). Last, the effects of *Euplotes sp*. body size on its own ecological dynamics and that of the *T. pyriformis* and bacteria were consistent across interpolation methods (Appendix II Figure 13-15 Eco-Pheno, Pheno-Eco and Pheno-Pheno panels). As CCM and related methods grow in use (Wang *et al*. 2016b; Matsuzaki *et al*. 2018; Barraquand *et al*. 2021; Doi *et al*. 2021; Kitayama *et al*. 2021), there is a real need for these tools to be robust even in the face of imperfect data, including missing timepoints. Our results therefore underline the needs to better understand how missing data may affect CCM and other time-series analyses and how to best predict missing data for analysis.

Last, because CCM analysis quantifies the effects of one time series on another but not on itself, we were unable measure density dependence within each species. Therefore, while our research showed the potential of using CCM in understanding complex casual effects between species ecological and phenotypic dynamics, we also notice the importance of combining time series analyses with other, possibly experimental, methods to measure causality for intra-specific interactions.

### Concluding remarks

Rapid phenotypic change has been suggested as a main driver of food web rewiring in future climates (Barbour & Gibert 2021), but little experimental evidence exists. Here, our results show that strong feedbacks between ecological and phenotypic dynamics depend on environmental conditions (temperature and nutrients), suggesting that rapid phenotypic change influences food web responses to environmental change. Moreover, the joint effects of temperature and nutrients do not equally affect all members of the community, as higher trophic levels are more likely to experience both independent and joint effects. Together, our results emphasize the need to incorporate phenotypic dynamics in future studies of food web responses to warming and eutrophication in a changing world and show how shifts in distinct environmental stressors can have complex but systematic effects on food web dynamics.

## ACKNOWLEDGMENTS

This work was supported by a U.S. Department of Energy, Office of Science, Office of Biological and Environmental Research, Genomic Science Program Grant under Award Number DE-SC0020362 to JPG.

