## Appendix for "Temperature and nutrients drive eco-phenotypic dynamics in a microbial food web"

### Supplementary Information: The interactive effects of temperature and nutrients on ecological and phenotypic dynamics across trophic levels

Ze-Yi Han[1], Daniel J. Wieczynski[1], Andrea Yammine[1], Jean P. Gibert[1]
[1]Department of Biology, Duke University, Durham, North Carolina, USA

**Index**

Appendix 1: Statistical Model Results ………………………………………………………. 1-3

Appendix 2: CCM Convergence Plot………………………………………………………… 4-17

#### Appendix I: Statistical modeling results

Table 1. GAMM models and model selection. In all models, we treated replicates as a random effect and day and treatment as fixed effects. We accounted for temporal autocorrelation using Autoregressive Moving Average (ARMA) correlation structure in the GAMM models using "corARMA" in R package 'nlme' (version 3.1-148).

| Species | Ecological or Phenotypic Dynamics | Models | Full 22°C smooth term significance | Full 25°C smooth term significance | Half 22°C smooth term significance | Half 25°C smooth term significance | AIC (lme) |
|---|---|---|---|---|---|---|---|
| *Bacteria* | Ecological Dynamics | OD ~ s(day, by = tre, k =10) + tre ,random = list(rep= ~ 1) | < 2e-16 *** | 3.57e-12 *** | 2.55e-05 *** | 0.0118 * | -1489.63 |
| | | OD ~ s(day, by = tre, k =10) + tre ,random = list(rep= ~ 1), correlation = corARMA(form = ~ day\|tre/rep, p = 1)) | < 2e-16 *** | 1.48e-13 *** | 8.12e-06 *** | 0.00991 ** | -1489.67 |
| | | **OD ~ s(day, by = tre, k =10) + tre,random = list(rep= ~ 1), correlation = corARMA(form = ~ day\|tre/rep, p = 2))** | **< 2e-16 ***** | **1.71e-11 ***** | **4.19e-06 ***** | **0.00935 **** | **-1499.45** |
| | | OD ~ s(day, by = tre, k =10) + tre ,random = list(rep= ~ 1), correlation = corARMA(form = ~ day\|tre/rep, p = 3)) | < 2e-16 *** | 1.36e-11 *** | 3.41e-06 *** | 0.00895 ** | -1497.48 |
| *Tetrahymena pyriformis* | Ecological Dynamics | log_density ~ s(day, by = tre, k =10) + tre, random = list(rep= ~ 1) | < 2e-16 *** | < 2e-16 *** | <2e-16 *** | < 2e-16 *** | 833.356 |
| | | **log_density ~ s(day, by = tre, k =10) + tre, random = list(rep= ~ 1), correlation = corARMA(form = ~ day\|tre/rep, p = 1)** | **< 2e-16 ***** | **< 2e-16 ***** | **< 2e-16 ***** | **< 2e-16 ***** | **779.954** |
| | | log_density ~ s(day, by = tre, k =10) + tre, random = list(rep= ~ 1), correlation = corARMA(form = ~ day\|tre/rep, p = 2) | < 2e-16 *** | < 2e-16 *** | < 2e-16 *** | < 2e-16 *** | 779.467 |
| | | log_density ~ s(day, by = tre, k =10) + tre, random = list(rep= ~ 1), correlation = corARMA(form = ~ day\|tre/rep, p = 3)) | < 2e-16 *** | < 2e-16 *** | < 2e-16 *** | < 2e-16 *** | 781.448 |
| *Euplotes sp.* | Ecological Dynamics | **log_density ~ s(day, by = tre, k =10) + tre, random = list(rep= ~ 1)** | **<2e-16 ***** | **<2e-16 ***** | **<2e-16 ***** | **<2e-16 ***** | **512.24** |
| | | log_density ~ s(day, by = tre, k =10) + tre, random = list(rep= ~ 1), correlation = corARMA(form = ~ day\|tre/rep, p = 1)) | <2e-16 *** | <2e-16 *** | <2e-16 *** | <2e-16 *** | 513.804 |
| | | log_density ~ s(day, by = tre, k =10) + tre, random = list(rep= ~ 1), correlation = corARMA(form = ~ day\|tre/rep, p = 2) | <2e-16 *** | <2e-16 *** | <2e-16 *** | <2e-16 *** | 514.193 |
| | | log_density ~ s(day, by = tre, k =10) + tre, random = list(rep= ~ 1), correlation = corARMA(form = ~ day\|tre/rep, p = 3) | <2e-16 *** | <2e-16 *** | <2e-16 *** | <2e-16 *** | 515.668 |
| *Tetrahymena pyriformis* | Phenotypic Dynamics | area ~ s(day, by = tre, k =7) + tre, random = list(rep= ~ 1) | <2e-16 *** | <2e-16 *** | <2e-16 *** | <2e-16 *** | 2139.42 |
| | | area ~ s(day, by = tre, k =7) + tre, random = list(rep= ~ 1), correlation = corARMA(form = ~ day\|tre/rep, p = 1) | <2e-16 *** | <2e-16 *** | <2e-16 *** | <2e-16 *** | 2131.18 |
| | | area ~ s(day, by = tre, k =7) + tre, random = list(rep= ~ 1), correlation = corARMA(form = ~ day\|tre/rep, p = 2) | <2e-16 *** | <2e-16 *** | <2e-16 *** | <2e-16 *** | 2132.75 |
| | | **area ~ s(day, by = tre, k =7) + tre, random = list(rep= ~ 1), correlation = corARMA(form = ~ day\|tre/rep, p = 3)** | **<2e-16 ***** | **<2e-16 ***** | **<2e-16 ***** | **<2e-16 ***** | **2126.74** |
| *Euplotes sp.* | Phenotypic Dynamics | area ~ s(day, by=tre, k = 9) + tre, random = list(rep= ~ 1) | 1.20e-13 *** | 3.84e-11 *** | 3.93e-15 *** | 6.20e-13 *** | 4559.91 |
| | | **area ~ s(day, by=tre, k = 9) + tre, random = list(rep= ~ 1), correlation = corARMA(form = ~ day\|tre/rep, p = 1)** | **9.03e-16 ***** | **3.04e-13 ***** | **< 2e-16 ***** | **5.91e-16 ***** | **4556.98** |
| | | area ~ s(day, by=tre, k = 9) + tre, random = list(rep= ~ 1), correlation = corARMA(form = ~ day\|tre/rep, p = 2) | 7.37e-16 *** | 3.47e-13 *** | < 2e-16 *** | 3.72e-16 *** | 4558.87 |
| | | area ~ s(day, by=tre, k = 9) + tre, random = list(rep= ~ 1), correlation = corARMA(form = ~ day\|tre/rep, p = 3) | 2.02e-15 *** | 1.76e-13 *** | < 2e-16 *** | < 2e-16 *** | 4560.77 |

Table 2. Quantifying temperature and nutrient effects on species temporal ecological and phenotypic dynamics using ARMA-GAMMs. Because we have two temperature and nutrients levels, we treated them as categorical variables in the models. We present model comparison using AICc on models that accounts for combinations of additive and/or interactive effects temperature and nutrient. Models are presented in the same order within each species for comparison.

<table>
<tr><th>Species</th><th>Ecological or Phenotypic Dynamics</th><th>Models</th><th>Temp effect estimates</th><th>Standard Error</th><th>p-value</th><th>Nutrient effect estimates</th><th>Stardard Error</th><th>p-value</th><th>Temp * Nut effect estimates</th><th>Stardard Error</th><th>p-value</th><th>22°C smooth term significance</th><th>25°C smooth term significance</th><th>Low Nutrient smooth term significance</th><th>High nutrient smooth term significance</th><th>AIC (lme)</th></tr>
<tr><td rowspan="8">Bacteria</td><td rowspan="8">Ecological Dynamics</td><td>OD ~ s(day, k = 10) + temp, random = list(all_rep = ~1), correlation = corARMA(form = ~ day|all_rep, p = 2)</td><td>-0.003025</td><td>0.002606</td><td>0.247</td><td>/</td><td>/</td><td>/</td><td>/</td><td>/</td><td>/</td><td>/</td><td>/</td><td>/</td><td>/</td><td>-1485.825</td></tr>
<tr><td>OD ~ s(day, k = 10) + nut,random = list(all_rep= ~ 1), correlation = corARMA(form = ~ day|all_rep, p = 2)</td><td>/</td><td>/</td><td>/</td><td>0.009648</td><td>0.001714</td><td>5.78e-08 ***</td><td>/</td><td>/</td><td>/</td><td>/</td><td>/</td><td>/</td><td>/</td><td>-1503.813</td></tr>
<tr><td>OD ~ s(day, k = 10) + nut + temp, random = list(all_rep= ~ 1), correlation = corARMA(form = ~ day|all_rep, p = 2)</td><td>-0.00328</td><td>0.001569</td><td>0.0377 *</td><td>0.009726</td><td>0.001569</td><td>2.97e-09 ***</td><td>/</td><td>/</td><td>/</td><td>/</td><td>/</td><td>/</td><td>/</td><td>-1505.806</td></tr>
<tr><td>OD ~ s(day, by = temp, k = 10) + temp , random = list(all_rep = ~1), correlation = corARMA(form = ~ day|all_rep, p = 2)</td><td>-0.003391</td><td>0.002681</td><td>0.207</td><td>/</td><td>/</td><td>/</td><td>/</td><td>/</td><td>/</td><td>3.72e-13 ***</td><td>3.56e-08 ***</td><td>/</td><td>/</td><td>-1475.198</td></tr>
<tr><td>OD ~ s(day, by = nut, k = 10) + nut , random = list(all_rep = ~1), correlation = corARMA(form = ~ day|all_rep, p = 2)</td><td>/</td><td>/</td><td>/</td><td>0.009779</td><td>0.001511</td><td>6.86e-10 ***</td><td>/</td><td>/</td><td>/</td><td>/</td><td>/</td><td>4.64e-08 ***</td><td>< 2e-16 ***</td><td>-1510.998</td></tr>
<tr><td>OD ~ s(day, by = temp, k = 10) + nut + temp, random = list(all_rep = ~1), correlation = corARMA(form = ~ day|all_rep, p = 2)</td><td>-0.00339</td><td>0.001603</td><td>0.0356 *</td><td>0.009738</td><td>0.001576</td><td>3.42e-09 ***</td><td>/</td><td>/</td><td>/</td><td>2.68e-15 ***</td><td>3.91e-09 ***</td><td>/</td><td>/</td><td>-1494.964</td></tr>
<tr><td>OD ~ s(day, by = nut, k = 10) + nut+ temp, random = list(all_rep = ~1), correlation = corARMA(form = ~ day|all_rep, p = 2)</td><td>-0.003394</td><td>0.001293</td><td>0.00931 **</td><td>0.009781</td><td>0.001296</td><td>1.42e-12 ***</td><td>/</td><td>/</td><td>/</td><td>/</td><td>/</td><td>4.28e-08 ***</td><td>< 2e-16 ***</td><td>-1514.961</td></tr>
<tr><td>OD ~ s(day, k = 10) + temp*nut, random = list(all_rep = ~1), correlation = corARMA(form = ~ day|all_rep, p = 2)</td><td>-0.005628</td><td>0.002138</td><td>0.009102 **</td><td>0.007449</td><td>0.002124</td><td>0.000555 ***</td><td>0.00463</td><td>0.003013</td><td>0.125897</td><td>/</td><td>/</td><td>/</td><td>/</td><td>-1506.116</td></tr>
<tr><td rowspan="8">Tetrahymena pyriformis</td><td rowspan="8">Ecological Dynamics</td><td>log_density ~ s(day, k = 10) + temp, random = list(all_rep = ~1), correlation = corARMA(form = ~ day|temp/all_rep, p = 1)</td><td>-1.0416</td><td>0.2642</td><td>0.000102 ***</td><td>/</td><td>/</td><td>/</td><td>/</td><td>/</td><td>/</td><td>/</td><td>/</td><td>/</td><td>/</td><td>899.707</td></tr>
<tr><td>log_density ~ s(day, k = 10) + nut,random = list(all_rep= ~ 1), correlation = corARMA(form = ~ day|nut/all_rep, p = 1)</td><td>/</td><td>/</td><td>/</td><td>0.3327</td><td>0.3017</td><td>0.271</td><td>/</td><td>/</td><td>/</td><td>/</td><td>/</td><td>/</td><td>/</td><td>911.337</td></tr>
<tr><td>log_density ~ s(day, k = 10) + nut + temp,random = list(all_rep= ~ 1), correlation = corARMA(form = ~ day|all_rep, p = 1)</td><td>-1.0478</td><td>0.2597</td><td>7.09e-05 ***</td><td>0.349</td><td>0.2597</td><td>0.18</td><td>/</td><td>/</td><td>/</td><td>/</td><td>/</td><td>/</td><td>/</td><td>899.927</td></tr>
<tr><td>log_density ~ s(day, by = temp, k = 10) + temp, random = list(all_rep = ~1), correlation = corARMA(form = ~ day|temp/all_rep, p = 1)</td><td>-1.2795</td><td>0.1629</td><td>9.58e-14 ***</td><td>/</td><td>/</td><td>/</td><td>/</td><td>/</td><td>/</td><td><2e-16 ***</td><td><2e-16 ***</td><td>/</td><td>/</td><td>781.2841</td></tr>
<tr><td>log_density ~ s(day, by = nut, k = 10) + nut, random = list(all_rep = ~1), correlation = corARMA(form = ~ day|nut/all_rep, p = 1)</td><td>/</td><td>/</td><td>/</td><td>0.4068</td><td>0.3044</td><td>0.183</td><td>/</td><td>/</td><td>/</td><td>/</td><td>/</td><td><2e-16 ***</td><td><2e-16 ***</td><td>930.061</td></tr>
<tr><td>log_density ~ s(day, by = temp, k = 10) + nut + temp , random = list(all_rep = ~1), correlation = corARMA(form = ~ day|all_rep, p = 1)</td><td>-1.2799</td><td>0.1555</td><td>8.28e-15 ***</td><td>0.3755</td><td>0.1528</td><td>0.0146 *</td><td>/</td><td>/</td><td>/</td><td><2e-16 ***</td><td><2e-16 ***</td><td>/</td><td>/</td><td>777.514</td></tr>
<tr><td>log_density ~ s(day, by = nut, k = 10) + temp+ nut, random = list(all_rep = ~1), correlation = corARMA(form = ~ day|all_rep, p = 1)</td><td>-1.0544</td><td>0.2521</td><td>3.9e-05 ***</td><td>0.4066</td><td>0.2585</td><td>0.117</td><td>/</td><td>/</td><td>/</td><td>/</td><td>/</td><td><2e-16 ***</td><td><2e-16 ***</td><td>917.817</td></tr>
<tr><td>log_density ~ s(day, k = 10) + temp*nut, random = list(all_rep = ~1), correlation = corARMA(form = ~ day|all_rep, p = 1)</td><td>-1.09252</td><td>0.36794</td><td>0.00325 **</td><td>0.30421</td><td>0.36794</td><td>0.40906</td><td>0.08948</td><td>0.52034</td><td>0.86359</td><td>/</td><td>/</td><td>/</td><td>/</td><td>901.8965</td></tr>
<tr><td rowspan="8">Euplotes sp.</td><td rowspan="8">Ecological Dynamics</td><td>log_density ~ s(day, k = 10) + temp, random = list(all_rep = ~1)</td><td>0.43257</td><td>0.06853</td><td>1.08e-09 ***</td><td>/</td><td>/</td><td>/</td><td>/</td><td>/</td><td>/</td><td>/</td><td>/</td><td>/</td><td>/</td><td>541.024</td></tr>
<tr><td>log_density ~ s(day, k = 10) + nut,random = list(all_rep= ~ 1)</td><td>/</td><td>/</td><td>/</td><td>-0.08742</td><td>0.10358</td><td>0.399</td><td>/</td><td>/</td><td>/</td><td>/</td><td>/</td><td>/</td><td>/</td><td>568.395</td></tr>
<tr><td>log_density ~ s(day, k = 10) + nut + temp,random = list(all_rep= ~ 1)</td><td>0.43257</td><td>0.06845</td><td>1.04e-09 ***</td><td>-0.08742</td><td>0.06845</td><td>0.203</td><td>/</td><td>/</td><td>/</td><td>/</td><td>/</td><td>/</td><td>/</td><td>541.381</td></tr>
<tr><td>log_density ~ s(day, by = temp, k = 10)+ temp , random = list(all_rep = ~1)</td><td>0.43257</td><td>0.06432</td><td>1.02e-10 ***</td><td>/</td><td>/</td><td>/</td><td>/</td><td>/</td><td>/</td><td><2e-16 ***</td><td><2e-16 ***</td><td>/</td><td>/</td><td>527.537</td></tr>
<tr><td>log_density ~ s(day, by = nut, k = 10) + nut , random = list(all_rep = ~1)</td><td>/</td><td>/</td><td>/</td><td>-0.08742</td><td>0.10358</td><td>0.399</td><td>/</td><td>/</td><td>/</td><td>/</td><td>/</td><td><2e-16 ***</td><td><2e-16 ***</td><td>571.787</td></tr>
<tr><td>log_density ~ s(day, by = temp, k = 10) + nut + temp, random = list(all_rep = ~1)</td><td>0.43257</td><td>0.06421</td><td>9.63e-11 ***</td><td>-0.08742</td><td>0.06421</td><td>0.175</td><td>/</td><td>/</td><td>/</td><td><2e-16 ***</td><td><2e-16 ***</td><td>/</td><td>/</td><td>527.6701</td></tr>
<tr><td>log_density ~ s(day, by = nut, k = 10) +nut + temp, random = list(all_rep = ~1)</td><td>0.43257</td><td>0.06639</td><td>3.48e-10 ***</td><td>-0.08742</td><td>0.06639</td><td>0.189</td><td>/</td><td>/</td><td>/</td><td>/</td><td>/</td><td><2e-16 ***</td><td><2e-16 ***</td><td>544.2109</td></tr>
<tr><td>log_density ~ s(day, k = 10) + temp*nut, random = list(all_rep = ~1)</td><td>0.33702</td><td>0.09663</td><td>0.000566 ***</td><td>-0.18298</td><td>0.09663</td><td>0.059323 .</td><td>0.19111</td><td>0.13665</td><td>0.163082</td><td>/</td><td>/</td><td>/</td><td>/</td><td>541.405</td></tr>
<tr><td rowspan="8">Tetrahymena pyriformis</td><td rowspan="8">Phenotypic Dynamics</td><td>area ~ s(day, k = 7) + temp, random = list(all_rep = ~1),correlation = corARMA(form = ~ day|temp/all_rep, p = 3)</td><td>-37.762</td><td>13.643</td><td>0.00626 **</td><td>/</td><td>/</td><td>/</td><td>/</td><td>/</td><td>/</td><td>/</td><td>/</td><td>/</td><td>/</td><td>2141.414</td></tr>
<tr><td>area ~ s(day, k = 7) + nut,random = list(all_rep= ~ 1), correlation = corARMA(form = ~ day|nut/all_rep, p = 3)</td><td>/</td><td>/</td><td>/</td><td>48.153</td><td>11.8</td><td>6.85e-05 ***</td><td>/</td><td>/</td><td>/</td><td>/</td><td>/</td><td>/</td><td>/</td><td>2135.699</td></tr>
<tr><td>area ~ s(day, k = 7) + nut + temp,random = list(all_rep= ~ 1), correlation = corARMA(form = ~ day|all_rep, p = 3)</td><td>-38.795</td><td>9.796</td><td>0.00011 ***</td><td>49.001</td><td>9.26</td><td>3.67e-07 ***</td><td>/</td><td>/</td><td>/</td><td>/</td><td>/</td><td>/</td><td>/</td><td>2125.098</td></tr>
<tr><td>area ~ s(day, by = temp, k = 7) + temp, random = list(all_rep = ~1), correlation = corARMA(form = ~ day|temp/all_rep, p = 3)</td><td>-48.219</td><td>22.899</td><td>0.0367 *</td><td>/</td><td>/</td><td>/</td><td>/</td><td>/</td><td>/</td><td><2e-16 ***</td><td><2e-16 ***</td><td>/</td><td>/</td><td>2152.360</td></tr>
<tr><td>area ~ s(day, by = nut, k = 7) + nut, random = list(all_rep = ~1), correlation = corARMA(form = ~ day|nut/all_rep, p = 3)</td><td>/</td><td>/</td><td>/</td><td>55.164</td><td>13.089</td><td>4.08e-05 ***</td><td>/</td><td>/</td><td>/</td><td>/</td><td>/</td><td><2e-16 ***</td><td><2e-16 ***</td><td>2131.108</td></tr>
<tr><td>area ~ s(day, by = temp, k = 7) + nut + temp, random = list(all_rep = ~1), correlation = corARMA(form = ~ day|all_rep, p = 3)</td><td>-45.377</td><td>20.821</td><td>0.0307 *</td><td>48.182</td><td>9.29</td><td>6.12e-07 ***</td><td>/</td><td>/</td><td>/</td><td><2e-16 ***</td><td><2e-16 ***</td><td>/</td><td>/</td><td>2136.618</td></tr>
<tr><td>area ~ s(day, by = nut, k = 7) + temp + nut, random = list(all_rep = ~1), correlation = corARMA(form = ~ day|all_rep, p =3)</td><td>56.549</td><td>9.929</td><td>5.44e-08 ***</td><td>-42.354</td><td>10.314</td><td>6.28e-05 ***</td><td>/</td><td>/</td><td>/</td><td>/</td><td>/</td><td><2e-16 ***</td><td><2e-16 ***</td><td>2119.911</td></tr>
<tr><td>area ~ s(day, k = 7) + temp * nut, random = list(all_rep = ~1), correlation = corARMA(form = ~ day|all_rep, p = 3)</td><td>-55.795</td><td>12.962</td><td>2.81e-05 ***</td><td>36.548</td><td>11.66</td><td>0.00203 **</td><td>32.309</td><td>17.733</td><td>0.07021 .</td><td>/</td><td>/</td><td>/</td><td>/</td><td>2124.072</td></tr>
<tr><td rowspan="8">Euplotes sp.</td><td rowspan="8">Phenotypic Dynamics</td><td>area ~ s(day, k = 9) + temp, random = list(all_rep = ~1),correlation = corARMA(form = ~ day|temp/all_rep, p = 1)</td><td>-566</td><td>143.9</td><td>0.000107 ***</td><td>/</td><td>/</td><td>/</td><td>/</td><td>/</td><td>/</td><td>/</td><td>/</td><td>/</td><td>/</td><td>4551.526</td></tr>
<tr><td>area ~ s(day, k = 9) + nut,random = list(all_rep= ~ 1), correlation = corARMA(form = ~ day|nut/all_rep, p = 1)</td><td>/</td><td>/</td><td>/</td><td>251.5</td><td>163.7</td><td>0.126</td><td>/</td><td>/</td><td>/</td><td>/</td><td>/</td><td>/</td><td>/</td><td>4562.876</td></tr>
<tr><td>area ~ s(day, k = 9) + nut + temp,random = list(all_rep= ~ 1), correlation = corARMA(form = ~ day|all_rep, p = 1)</td><td>-569.7</td><td>142.6</td><td>8.42e-05 ***</td><td>252.9</td><td>142.6</td><td>0.0772 .</td><td>/</td><td>/</td><td>/</td><td>/</td><td>/</td><td>/</td><td>/</td><td>4550.373</td></tr>
<tr><td>area ~ s(day, by = temp, k = 9) + temp, random = list(all_rep = ~1), correlation = corARMA(form = ~ day|temp/all_rep, p = 1)</td><td>-569.84</td><td>131.7</td><td>2.17e-05 ***</td><td>/</td><td>/</td><td>/</td><td>/</td><td>/</td><td>/</td><td><2e-16 ***</td><td><2e-16 ***</td><td>/</td><td>/</td><td>4544.981</td></tr>
<tr><td>area ~ s(day, by = nut, k = 9) + nut, random = list(all_rep = ~1), correlation = corARMA(form = ~ day|nut/all_rep, p = 1)</td><td>/</td><td>/</td><td>/</td><td>244.8</td><td>163.8</td><td>0.136</td><td>/</td><td>/</td><td>/</td><td>/</td><td>/</td><td><2e-16 ***</td><td><2e-16 ***</td><td>4569.363</td></tr>
<tr><td>area ~ s(day, by = temp, k = 9) + nut + temp, random = list(all_rep = ~1), correlation = corARMA(form = ~ day|all_rep, p = 1)</td><td>-573.2</td><td>130.7</td><td>1.69e-05 ***</td><td>237.7</td><td>130.9</td><td>0.0705 .</td><td>/</td><td>/</td><td>/</td><td><2e-16 ***</td><td><2e-16 ***</td><td>/</td><td>/</td><td>4543.674</td></tr>
<tr><td>area ~ s(day, by = nut, k = 9) + temp + nut, random = list(all_rep = ~1), correlation = corARMA(form = ~ day|all_rep, p = 1)</td><td>-583</td><td>132.2</td><td>1.53e-05 ***</td><td>243.8</td><td>132</td><td>0.0659 .</td><td>/</td><td>/</td><td>/</td><td>/</td><td>/</td><td><2e-16 ***</td><td><2e-16 ***</td><td>4554.992</td></tr>
<tr><td>area ~ s(day, k = 9) + temp *nut, random = list(all_rep = ~1), correlation = corARMA(form = ~ day|all_rep, p = 1)</td><td>-545.1</td><td>200.91</td><td>0.00712 **</td><td>278.24</td><td>203.91</td><td>0.17361</td><td>-49.82</td><td>286</td><td>0.86184</td><td>/</td><td>/</td><td>/</td><td>/</td><td>4552.342</td></tr>
</table>

Table 3. Linear models for temperature and nutrients effects on multiple aspect of ecological dynamics. Most parsimonious models are selected by AICc and are bolded. Significant additive and interactive effects of temperature and nutrients from the best models are highlighted and shown in main text Fig. 2. Models are listed in the same order within each species for easy comparison.

| Species | Descriptor | Models | Temperature effects Estimates | Standard Error | p-value | Nutrient effects Estimates | Standard Error | p-value | Temp * nutrient effect | Standard Error | p-value | R square | AIC |
|---|---|---|---|---|---|---|---|---|---|---|---|---|---|
| ***Bacteria*** | Initial population growth (IG) | b.IG ~ temp | -0.000175 | 0.0003895 | 0.659 | / | / | / | / | / | / | 0.01109 | -221.1834 |
| | | **b.IG ~ nutr** | / | / | / | 0.00045 | 0.0003771 | 0.248 | / | / | / | 0.07332 | **-222.4833** |
| | | b.IG ~ temp + nutr | -0.000175 | 0.0003857 | 0.656 | 0.00045 | 0.0003857 | 0.259 | / | / | / | 0.08441 | -220.7241 |
| | | b.IG ~ temp * nutr | 0.000125 | 0.0005521 | 0.823759 | 0.00075 | 0.0005521 | 0.193187 | -0.0006 | 0.0007808 | 0.453432 | 0.117 | -219.4488 |
| | Max Optimal Density (MOD) | MOD ~ temp | -0.00725 | 0.003971 | 0.0846. | / | / | / | / | / | / | 0.1562 | -128.3072 |
| | | MOD ~ nutr | / | / | / | 0.01425 | 0.002722 | 5.60e-05 *** | / | / | / | 0.6035 | -143.4138 |
| | | **MOD ~ temp + nutr** | -0.00725 | 0.00218 | 0.00401 ** | 0.01425 | 0.00218 | 5.090e-06 *** | / | / | / | 0.7598 | **-151.4334** |
| | | MOD ~ temp * nutr | -0.0095 | 0.003077 | 0.00613 ** | 0.011 | 0.003077 | 1.6e-05 *** | 0.0045 | 0.004352 | 0.31651 | 0.7748 | -150.7271 |
| | Coefficient of Variation (CV) | b.CV ~ temp | -0.00749 | 0.05396 | 0.891 | / | / | / | / | / | / | 0.001069 | -23.94258 |
| | | **b.CV ~ nutr** | / | / | / | 0.17521 | 0.03477 | 8.54e-05 *** | / | / | / | 0.5851 | **-41.51658** |
| | | b.CV ~ temp + nutr | -0.00749 | 0.03574 | 0.836474 | 0.17521 | 0.03574 | 0.000134 *** | / | / | / | 0.5862 | -39.5682 |
| | | b.CV ~ temp * nutr | -0.0072838 | 0.0520933 | 0.89055 | 0.1754129 | 0.0520933 | 0.00392 ** | -0.000413 | 0.0736711 | 0.9956 | 0.5862 | -37.56823 |
| ***Tetrahymena pyriformis*** | Initial population growth | **t.IG ~ temp** | 0.50026 | 0.07549 | 1.16e-06 *** | / | / | / | / | / | / | 0.6662 | **-8.996006** |
| | | t.IG ~ nutr | / | / | / | -0.039 | 0.1304 | 0.768 | / | / | / | 0.004049 | 17.241444 |
| | | t.IG ~ temp + nutr | 0.50026 | 0.0768 | 1.88e-06 *** | -0.039 | 0.0768 | 0.617 | / | / | / | 0.6703 | -7.288963 |
| | | t.IG ~ temp * nutr | 0.63996 | 0.10215 | 4.07e-06 *** | 0.1007 | 0.10215 | 0.336 | -0.2794 | 0.14446 | 0.0674 . | 0.7222 | -9.404116 |
| | Maximum Abundance (MA) | t.MA ~ temp | -212 | 479.5 | 0.663 | / | / | / | / | / | / | 0.008807 | 411.3144 |
| | | t.MA ~ nutr | / | / | / | 1765.5 | 300.5 | 6.51e-06 *** | / | / | / | 0.6108 | 388.8788 |
| | | t.MA ~ temp + nutr | -212 | 304 | 0.493 | 1765.5 | 304 | 9.17e-06 *** | / | / | / | 0.6196 | 390.3295 |
| | | **t.MA ~ temp * nutr** | -975.3 | 368.6 | 0.0155 * | 1002.2 | 368.6 | 0.0132 * | 1526.7 | 521.3 | 0.0083 ** | 0.7338 | **383.7636** |
| | Coefficient of Variation (CV) | t.CV ~ temp | 0.29779 | 0.05183 | 8.86e-06 *** | / | / | / | / | / | / | 0.6001 | -27.044711 |
| | | t.CV ~ nutr | / | / | / | -0.0898 | 0.07969 | 0.272 | / | / | / | 0.05457 | -6.396766 |
| | | t.CV ~ temp + nutr | 0.29779 | 0.0493 | 5.39e-06 *** | -0.0898 | 0.0493 | 0.0828 . | / | / | / | 0.6546 | -28.565539 |
| | | **t.CV ~ temp * nutr** | 0.20033 | 0.06445 | 0.00554 ** | -0.18727 | 0.06445 | 0.00875 ** | 0.19492 | 0.09115 | 0.04500 * | 0.7189 | **-31.507774** |
| | Time of Collapse (TC) | **TC ~ temp** | -2.7576 | 0.4954 | 1.59e-05 *** | / | / | / | / | / | / | 0.5961 | **77.05477** |
| | | TC ~ nutr | / | / | / | 0.7273 | 0.7631 | 0.351 | / | / | / | 0.04146 | 96.92986 |
| | | TC ~ temp + nutr | -2.7302 | 0.4899 | 1.87e-05 *** | 0.6032 | 0.4899 | 0.233 | / | / | / | 0.6245 | 77.37444 |
| | | TC ~ temp * nutr | -2.4333 | 0.7223 | 0.00322 ** | 0.9 | 0.7223 | 0.22789 | -0.5667 | 0.998 | 0.57681 | 0.6308 | 78.98743 |
| ***Euplotes sp.*** | Initial population growth | e.IG ~ temp | 0.09146 | 0.02018 | 0.000165 *** | / | / | / | / | / | / | 0.4828 | -72.31915 |
| | | e.IG ~ nutr | / | / | / | -0.02389 | 0.0276 | 0.396 | / | / | / | 0.03295 | -57.30049 |
| | | e.IG ~ temp + nutr | 0.09146 | 0.01999 | 0.000164 *** | -0.02389 | 0.01999 | 0.245279 | / | / | / | 0.5157 | -71.89895 |
| | | **e.IG ~ temp * nutr** | 0.02835 | 0.02099 | 0.191993 | -0.087 | 0.02099 | 0.000502 *** | 0.12621 | 0.02969 | 0.000391 *** | 0.7456 | **-85.34791** |
| | Maximum Abundance (MA) | e.MA ~ temp | 39.83 | 20.83 | 0.069 . | / | / | / | / | / | / | 0.1425 | 260.7729 |
| | | e.MA ~ nutr | / | / | / | 60.67 | 18.41 | 0.00329 ** | / | / | / | 0.3306 | 254.8308 |
| | | **e.MA ~ temp + nutr** | 39.83 | 16.71 | 0.02668 * | 60.67 | 16.71 | 0.00157 ** | / | / | / | 0.4731 | **251.0854** |
| | | e.MA ~ temp * nutr | 34.5 | 24.16 | 0.169 | 55.33 | 24.16 | 0.033 * | 10.67 | 0.758 | 0.758 | 0.4756 | 252.9687 |
| | Coefficient of Variation (CV) | e.CV ~ temp | -0.13712 | 0.03162 | 0.000265 *** | / | / | / | / | / | / | 0.4608 | -50.76312 |
| | | e.CV ~ nutr | / | / | / | 0.0983 | 0.03762 | 0.0159 * | / | / | / | 0.2368 | -42.42559 |
| | | e.CV ~ temp + nutr | -0.13712 | 0.02424 | 1.29e-05 *** | 0.0983 | 0.02424 | 0.000568 *** | / | / | / | 0.6977 | -62.647 |
| | | **e.CV ~ temp * nutr** | -0.07689 | 0.02951 | 0.01692 * | 0.15853 | 0.02951 | 2.94e-05 *** | -0.12045 | 0.04173 | 0.00913 ** | 0.7866 | **-69.00361** |
| | Time of Peak (TP) | **TP ~ temp** | -2.1667 | 0.8609 | 0.0196 * | / | / | / | / | / | / | 0.2235 | **107.8341** |
| | | TP ~ nutr | / | / | / | 1.3333 | 0.9347 | 0.168 | / | / | / | 0.08466 | 111.7836 |
| | | TP ~ temp + nutr | -2.1667 | 0.8317 | 0.0165 * | 1.3333 | 0.8317 | 0.1239 | / | / | / | 0.3082 | 107.0635 |
| | | TP ~ temp * nutr | -1.6667 | 1.1949 | 0.178 | 1.8333 | 1.1949 | 0.141 | -1 | 1.6898 | 0.561 | 0.3201 | 108.6469 |

**Appendix 2: CCM Convergence Plots**

1. Interpolating missing datapoints

Four datapoints were missing from our sampling regime corresponding to days 5-6 and 12-13 of the time series. To account for those missing datapoints for CCM analysis, we interpolated each time series of the ecological data and phenotypic data using three different methods: 1) linear interpolation, 2) spline interpolation, and 3) smooth spline interpolation using the 'approx', 'spline', and 'smooth.spline' functions in base R respectively. Because we do not know which of the interpolation methods yields predictions that are the closest to the true (unobserved) dynamics, we used all three interpolation variants (dubbed "linear", "spline", and "smooth spline") for the CCM analysis.

For each variant of the time series, we applied multispacial CCM analysis (using R package 'multispatialCCM' version 1.0) to quantify the strength of causal effects between species ecological dynamics and phenotypic dynamics. To better infer the causal effects within the system, we did not take into account CCM results that did not show clear convergence (thus no causality detected) based on Fig 1-12 in current appendix. To account for the differences between interpolation methods, we averaged the multispacial CCM values that showed causality (Fig. 13-15 in current appendix) across three sets of CCM results to generate the final results used in our inference (main text Fig. 4). However, as we show in the following section, the results of the CCM analysis done in each of the three datasets show only minor quantitative differences.

2. Multispacial CCM results for each of the three datasets

*2.1 Linear interpolation*

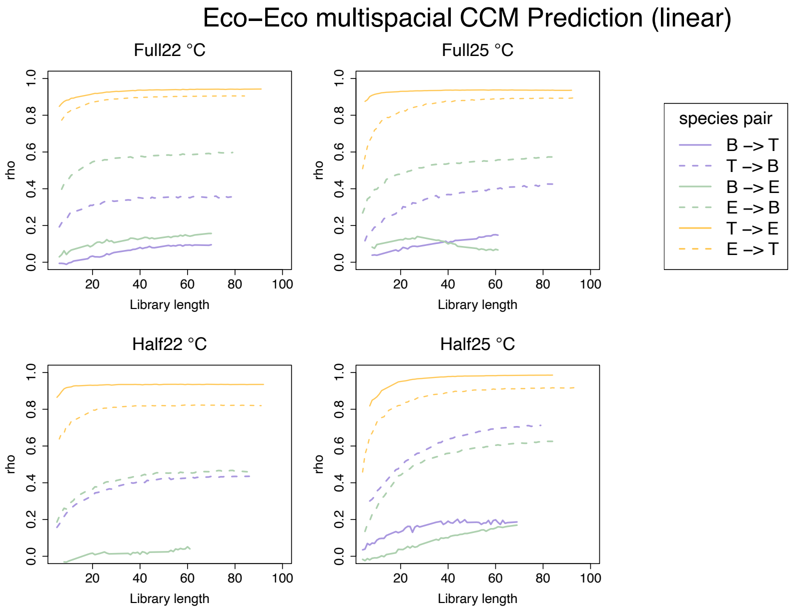


Figure S1. Eco-Eco effects in all treatments generated by multispacial CCM with linear interpolation of the missing data. CCM skill/ predictability of interactions between the ecological dynamics of Bacteria, *T. pyriformis* and *Euplotes sp.* and the phenotypic dynamics of *T. pyriformis* and *Euplotes sp.*. Due to the low counts for phenotypic data gathered near *T. pyriformis* population collapse time, the phenotypic data become unreliable. Therefore, we only used day 0-9 of the *T. pyriformis* phenotypic dynamics data in multispacial CCM test for estimating the species interactions. Because multispacial CCM require equal length of the time series, we also used day 0-9 of the rest of time series when they are paired with *T. pyriformis* phenotypic dynamics in multispacial CCM test. In the figure, "B -> T" denotes the bacteria ecological dynamics driving *T. pyriformis* ecological dynamics. Because the effects of bacteria ecological dynamics on *Euplotes sp.* "B -> E" (green solid line) in 25°C full nutrient level treatment does not show convergence (hence no causal effects detected), we did not use the CCM estimation skill (rho) for Fig 4 - 5.

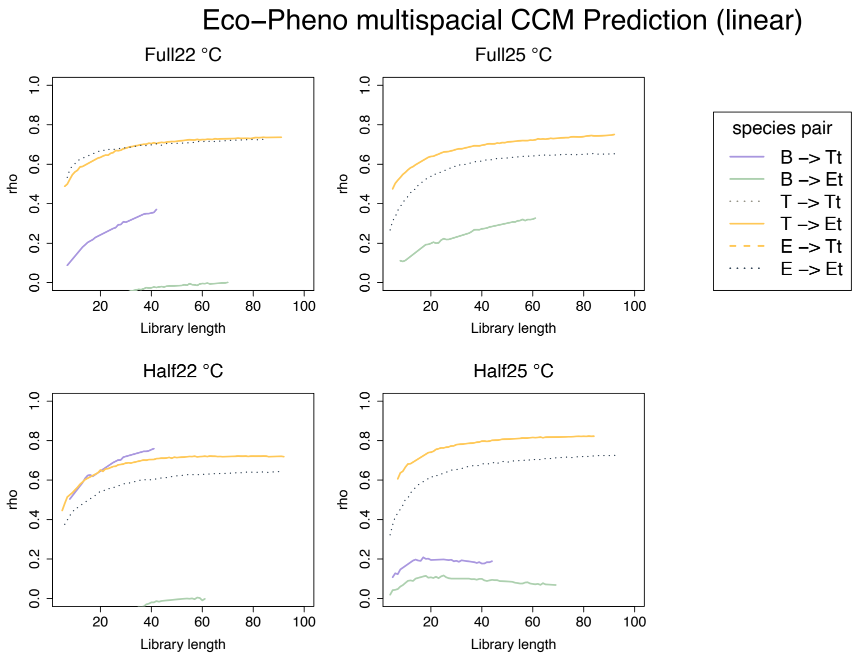


Figure S2. Eco-Pheno effects in all treatment generated by CCM with linear approximation. In the figure, “B -> Tt” denotes the bacteria ecological dynamics driving *T. pyriformis* phenotypic dynamics (trait changes).

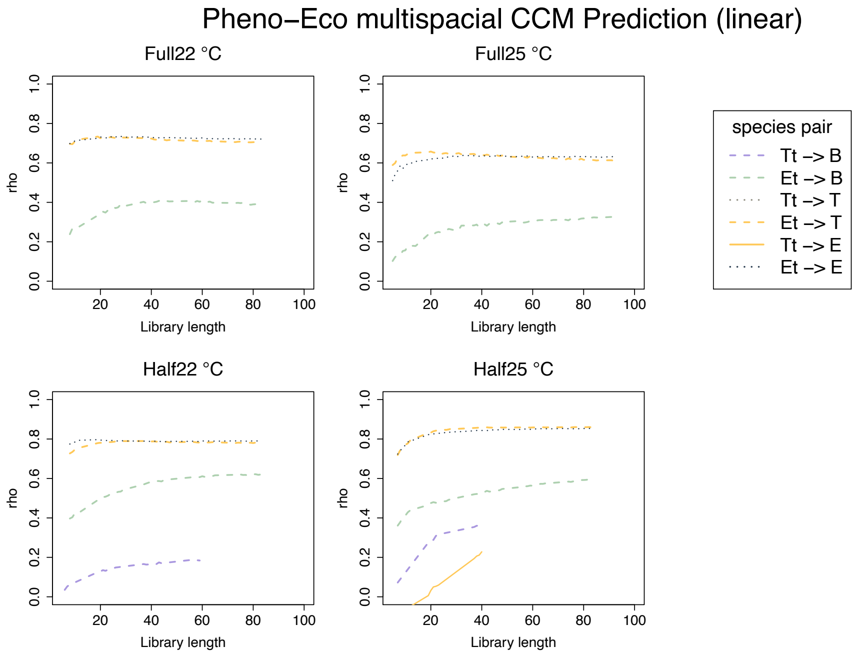


Figure S3. Pheno-Eco effects in all treatment generated by CCM with linear approximation.

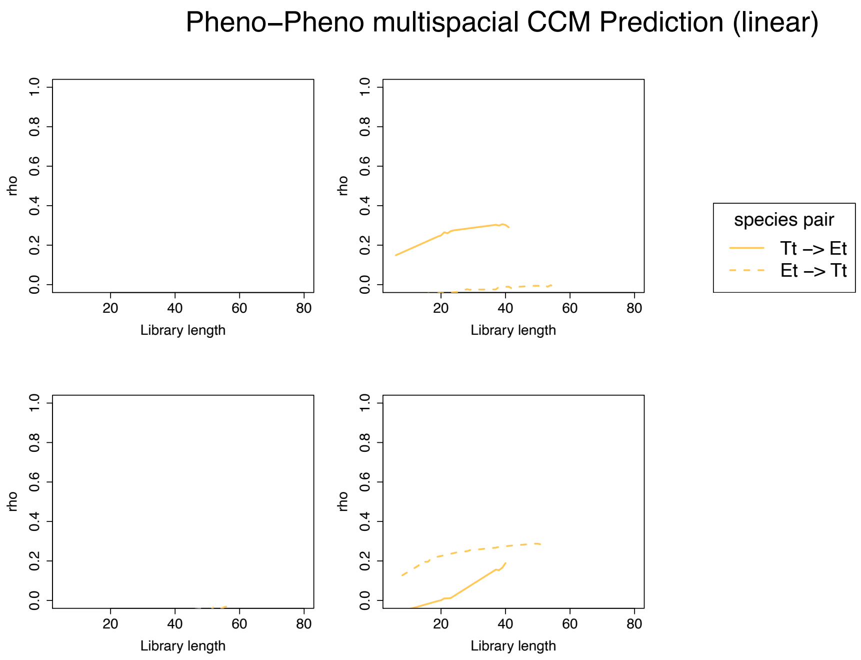


Figure S4. Pheno-Pheno effects in all treatment generated by CCM with linear approximation.

#### *2.2 Spline interpolation*

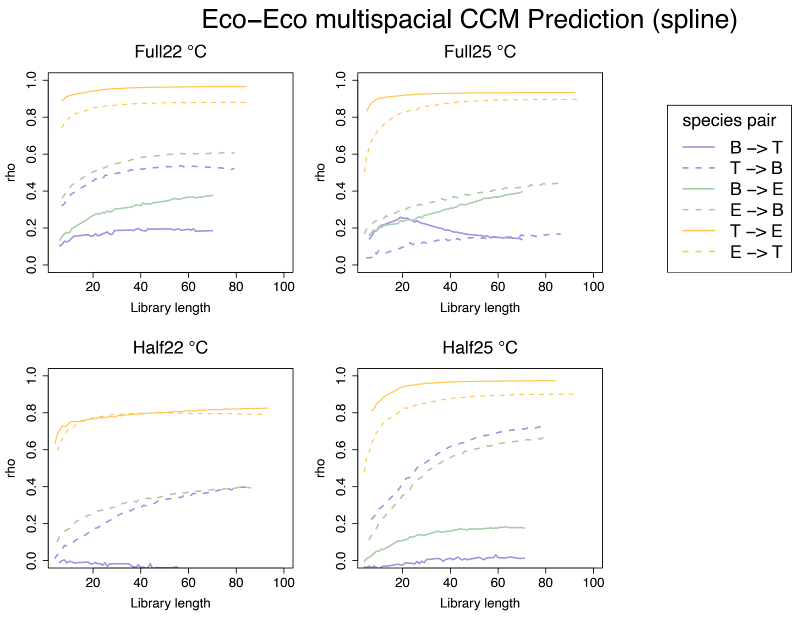


Figure S5. Eco-Eco effects in all treatment generated by CCM with spline interpolation. Because the effects of bacteria ecological dynamics on *Euplotes sp.* “B -> T” (purple solid line) in 25°C full nutrient level treatment does not show convergence (hence no causal effects detected), we did not use the CCM estimation skill (rho) for Fig 4 - 5.

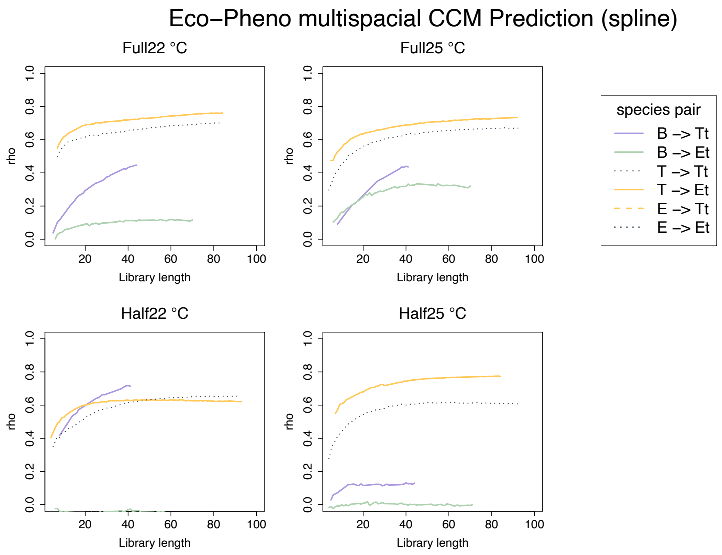


Figure S6. Eco-Pheno effects in all treatment generated by CCM with spline interpolation. Because the effects of bacteria ecological dynamics on *Euplotes sp.* phenotypic dynamics “B -> Et” (green solid line) in 25°C half nutrient level treatment does not show convergence (hence no causal effects detected), we did not use the CCM estimation skill (rho) for Fig 4 - 5.

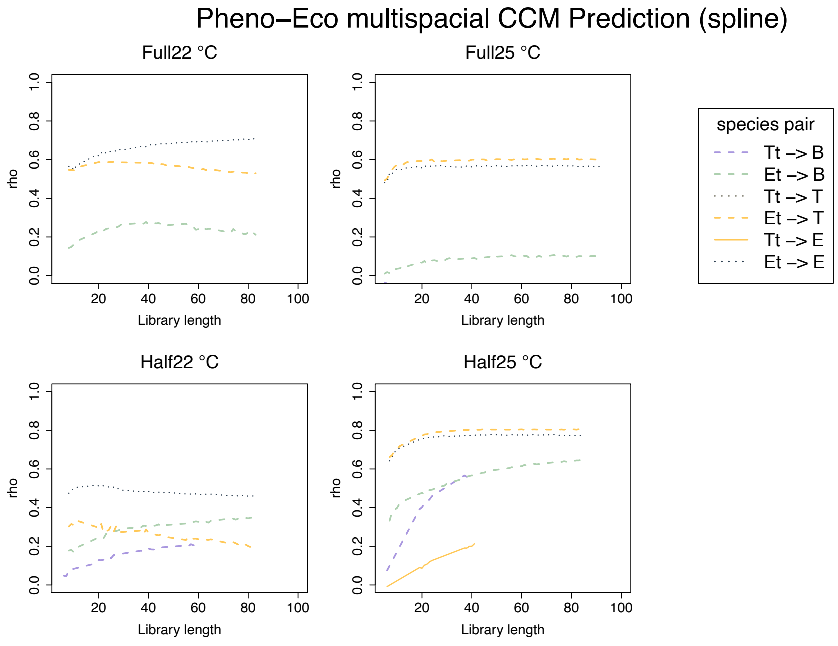


Figure S7. Pheno-Eco effects in all treatment generated by CCM with spline interpolation. Because the effects of *Euplotes sp.* phenotypic dynamics on *T. pyriformis* ecological dynamics “Et -> T” (yellow dash line) in 22°C half and full nutrient level treatments do not show convergence (hence no causal effects detected), we did not use the CCM estimation skill (rho) for Fig 4 - 5.

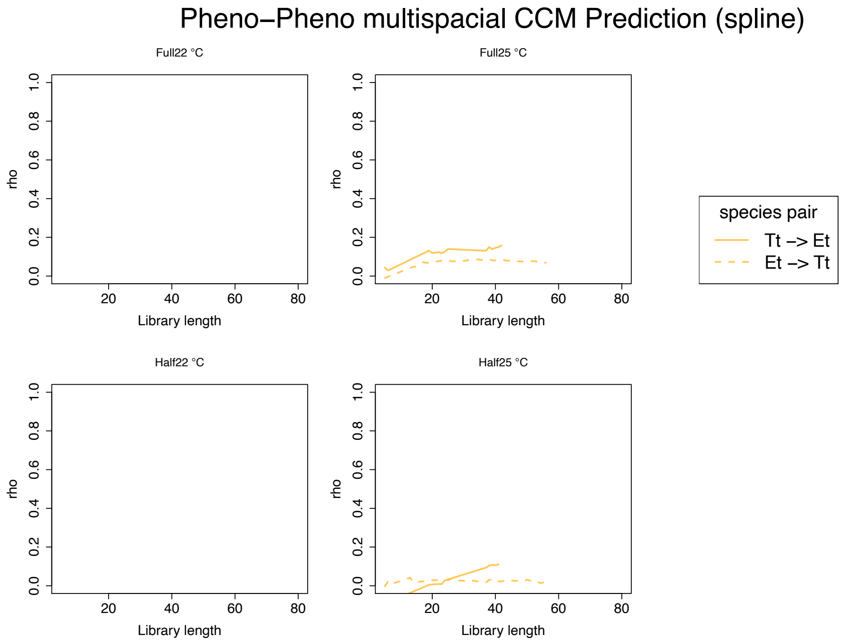


Figure S8. Pheno-Pheno effects in all treatment generated by CCM with spline interpolation. Because the effects of *Euplotes sp.* phenotypic dynamics on *T. pyriformis* phenotypic dynamics “Et -> Tt” (yellow dash line) in 25°C half nutrient level treatments do not show convergence (hence no causal effects detected), we did not use the CCM estimation skill (rho) for Fig 4 - 5.

*2.3 Smooth spline interpolation*

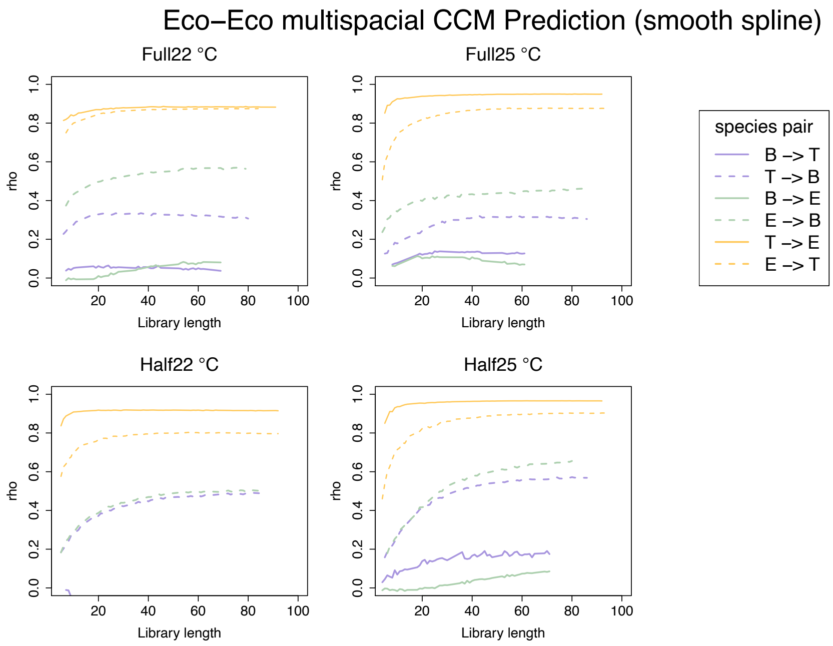


Figure S9. Eco-Eco effects in all treatment generated by CCM with smooth spline interpolation. Because the effects of bacteria ecological dynamics on bacteria ecological dynamics “B -> T” (purple solid line) in 22°C Full nutrient level treatment does not show convergence (hence no causal effects detected), we did not use the CCM estimation skill (rho) for Fig 4 - 5.

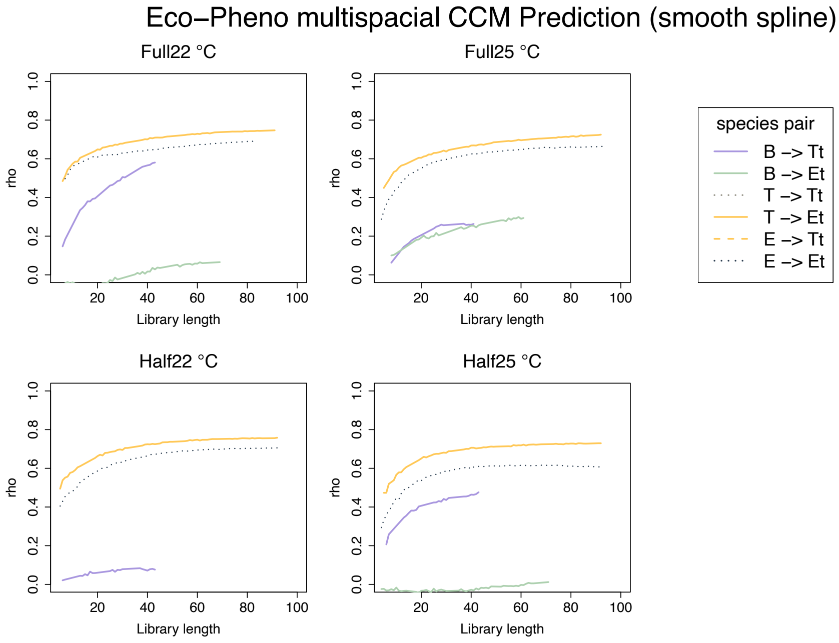


Figure S10. Eco-Pheno effects in all treatment generated by CCM with smooth spline interpolation. Because the effects of bacteria ecological dynamics on *Euplotes sp.* ecological dynamics “B -> E” (green solid line) in 25°C half nutrient level treatment do not show convergence (hence no causal effects detected), we did not use the CCM estimation skill (rho) for Fig 4 - 5.

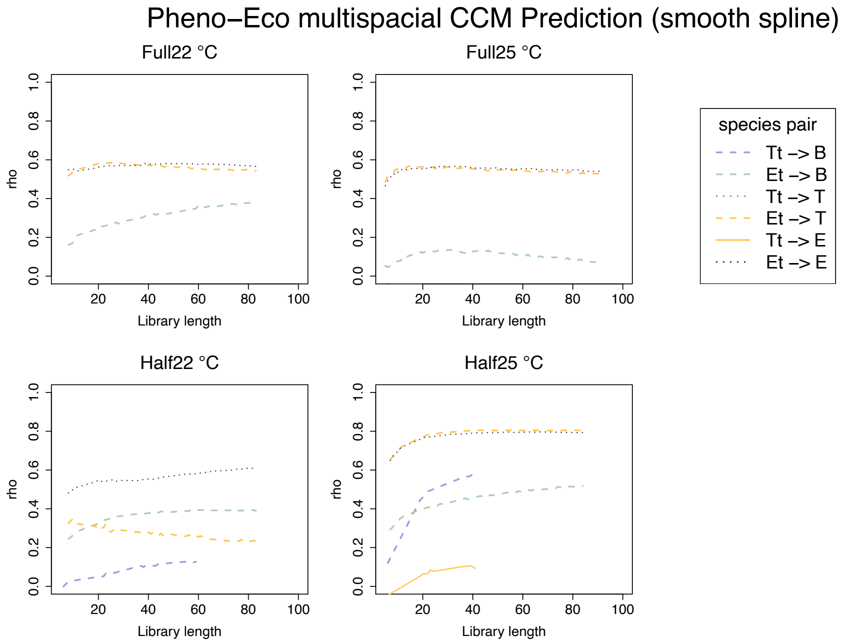


Figure S11. Pheno-Eco effects in all treatment generated by CCM with smooth spline interpolation. Because the effects of *Euplotes sp.* phenotypic dynamics on *T. pyriformis* ecological dynamics "Et -> T" (yellow dash line) in 22°C half nutrient level treatments do not show convergence (hence no causal effects detected), we did not use the CCM estimation skill (rho) for Fig 4 - 5.

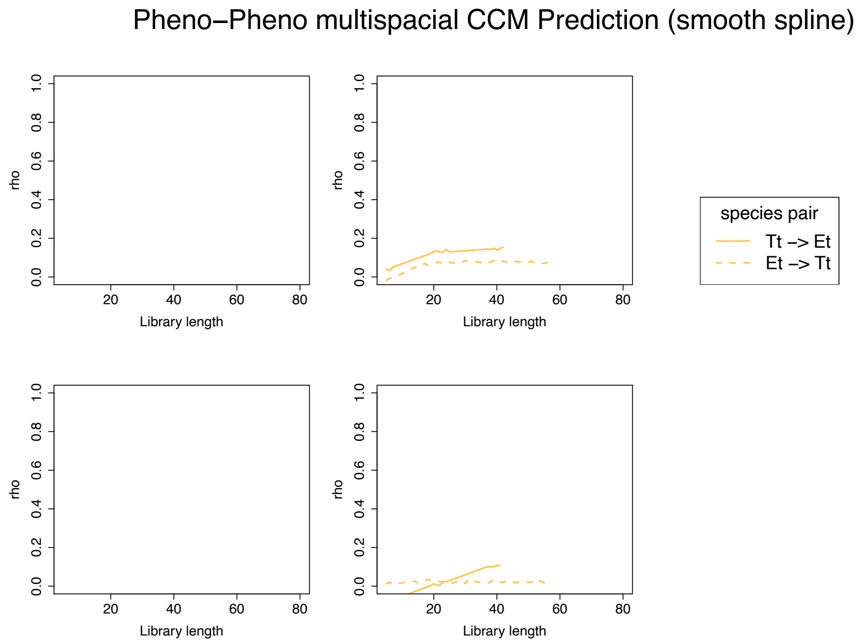


Figure S12. Pheno-Pheno effects in all treatment generated by CCM with smooth spline interpolation. Because the effects of *Euplotes sp.* phenotypic dynamics on *T. pyriformis* phenotypic dynamics "Et -> Tt" (yellow dash line) in 25°C half nutrient level treatments do not show convergence (hence no causal effects detected), we did not use the CCM estimation skill (rho) for Fig 4 - 5.

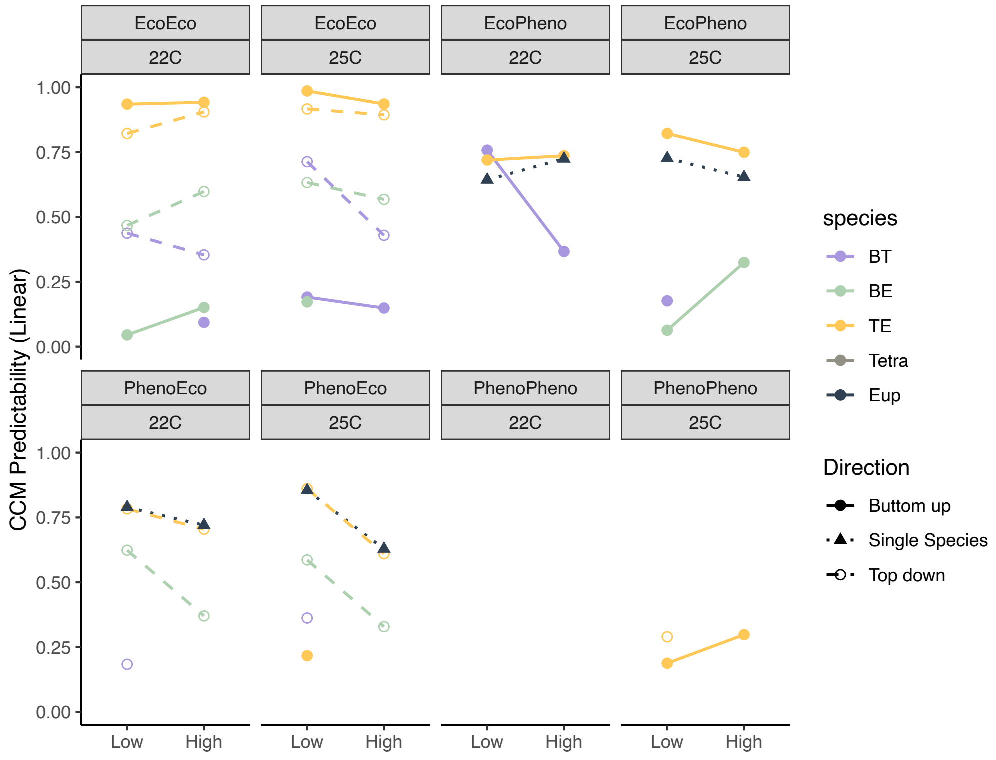


Figure S13. CCM predictabilities of ecological and phenotypic dynamics with linear interpolation across treatments for each species pair.

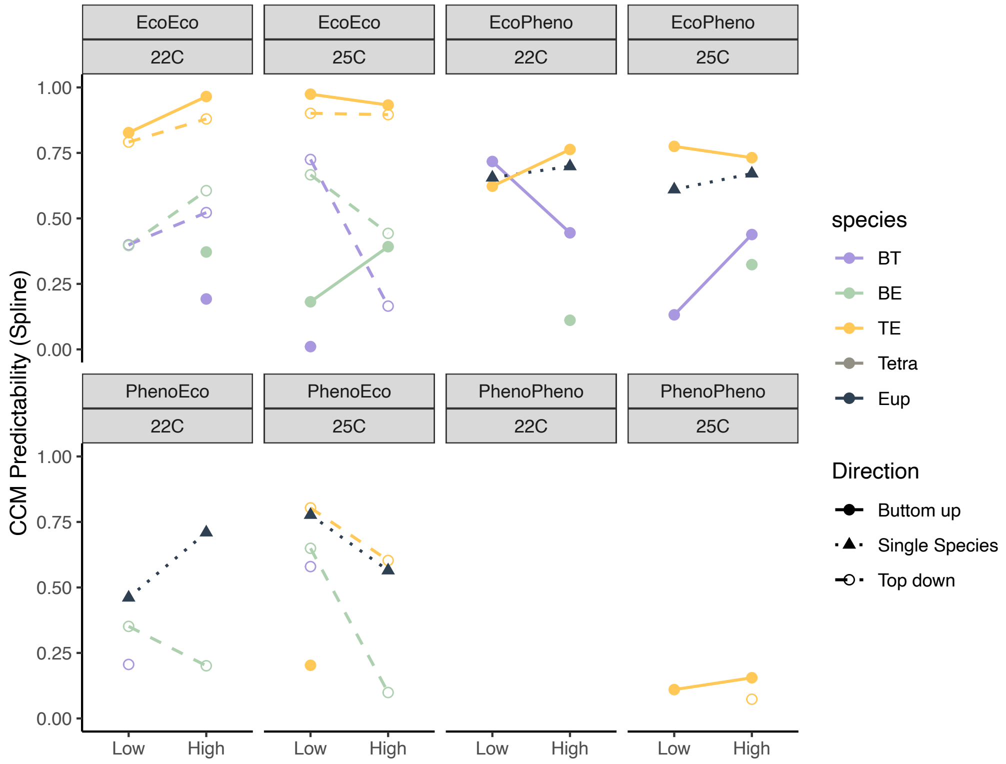

Figure S14. CCM predictabilities of ecological and phenotypic dynamics with spline interpolation across treatments for each species pair.

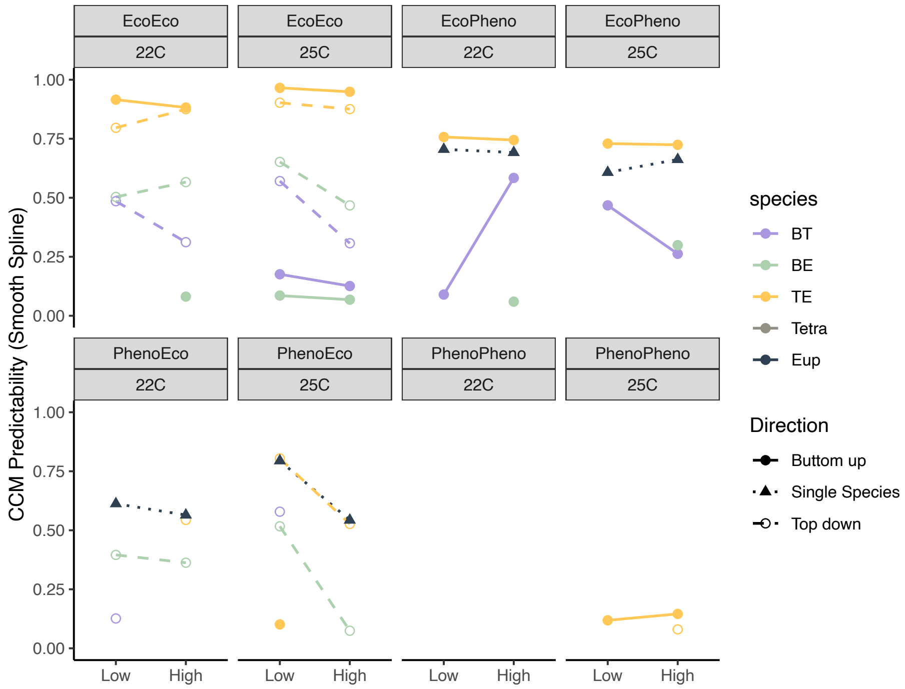


Figure S15. CCM predictabilities of ecological and phenotypic dynamics with spline interpolation across treatments for each species pair.
